# Prediction Under Uncertainty: Dissociating Sensory from Cognitive Expectations in Highly Uncertain Musical Contexts

**DOI:** 10.1101/2021.08.18.456640

**Authors:** Iris Mencke, David Ricardo Quiroga-Martinez, Diana Omigie, Georgios Michalareas, Franz Schwarzacher, Niels Trusbak Haumann, Peter Vuust, Elvira Brattico

**Affiliations:** Department of Music, Max Planck Institute for Empirical Aesthetics, Grüneburgweg 14, 60322 Frankfurt/Main, Germany; Department of Neuroscience, Max Planck Institute for Empirical Aesthetics, Grüneburgweg 14, 60322 Frankfurt/Main, Germany; Center for Music in the Brain, Department of Clinical Medicine, Aarhus University & The Royal Academy of Music, Aarhus/Aalborg, Nørrebrogade 44, 8000 Aarhus C, Denmark; Department of Psychology, Goldsmiths, SE14 6NW, University of London, London, United Kingdom; Department of Education, Psychology and Communication, University of Bari Aldo Moro, Piazza Umberto I, 70121 Bari, Italy

**Keywords:** predictive processing, MMN, tonal hierarchy, expectation, atonal music, MEG

## Abstract

Predictive models in the brain rely on the continuous extraction of regularities from the environment. These models are thought to be updated by novel information, as reflected in prediction error responses such as the mismatch negativity (MMN). However, although in real life individuals often face situations in which uncertainty prevails, it remains unclear whether and how predictive models emerge in high-uncertainty contexts. Recent research suggests that uncertainty affects the magnitude of MMN responses in the context of music listening. However, musical predictions are typically studied with MMN stimulation paradigms based on Western tonal music, which are characterized by relatively high predictability. Hence, we developed an MMN paradigm to investigate how the high uncertainty of atonal music modulates predictive processes as indexed by the MMN and behavior. Using MEG in a group of 20 subjects without musical training, we demonstrate that the magnetic MMN in response to pitch, intensity, timbre, and location deviants is evoked in both tonal and atonal melodies, with no significant differences between conditions. In contrast, in a separate behavioral experiment involving 39 non-musicians, participants detected pitch deviants more accurately and rated confidence higher in the tonal than in the atonal musical context. These results indicate that contextual tonal uncertainty modulates processing stages in which conscious awareness is involved, although deviants robustly elicit low-level pre-attentive responses such as the MMN. The achievement of robust MMN responses, despite high tonal uncertainty, is relevant for future studies comparing groups of listeners’ MMN responses to increasingly ecological music stimuli.

## 1. Introduction

Perception is increasingly seen as an active process by which the brain creates generative models of the environment in order to predict upcoming stimuli (Clark, 2013; Friston and Kiebel, 2009). When predictions are violated, prediction error responses arise, which are thought to index the update of predictive models by prediction error (Friston, 2005). However, in real life we often face and even derive pleasure from complex perceptual environments in which predictive models are difficult to establish and uncertainty prevails. In this study, we employ a musical mismatch negativity (MMN) paradigm comprising high- and low-uncertainty conditions as well as a related behavioral task in order to understand whether and how predictive models arise in such high-uncertainty perceptual environments.

Predictive models are formed by the regularity extracted from the preceding auditory sequence (Denham and Winkler, 2006) and the MMN is a brain response that is thought to reflect the continuous updating of such predictive models by prediction errors—i.e., the difference between expected and actual sensory inputs (Bendixen et al., 2012; Lieder et al., 2013; Näätänen, 2003; Rohrmeier and Koelsch, 2012; Vuust and Frith, 2008). The MMN responds to deviant sounds introduced in an otherwise regular sound sequence, indexes low-level auditory regularities (Koelsch, 2012; Näätänen, 1992) and serves as a marker of local deviance processing. Being an early and highly automatic response, it is regarded as a neural correlate of sensory expectations for basic sound features. Cognitive expectations, on the other hand, correspond to stylistic regularities such as tonal hierarchies or specific pitch patterns and are likely reflected in later components such as the P300, an evoked brain response occurring significantly later than the MMN (Polich, 2007). The P300 indexes the violation of global auditory expectations emerging from the context previously heard, that engage higher level processes in the brain (Bekinschtein et al., 2009; Wacongne et al., 2012) in which conscious awareness is engaged.

While early research used simple, short auditory patterns as stimuli (Fujioka et al., 2004; Tervaniemi et al., 2001; Zuijen et al., 2004), in more recent research musical MMN paradigms have introduced musical stimuli with higher complexity with regard to metrical, rhythmical or melodic aspects that allow for the study of contexts with varying degrees of predictability (Brattico et al., 2006; Huotilainen et al., 2009; Lumaca et al., 2018; Putkinen et al., 2014; Tervaniemi et al., 2014; Vuust et al., 2012). Some of these studies have shown that the MMNm is attenuated in melodic contexts with high uncertainty, especially for pitch deviants, and that behavioral measures of deviance detection and subjective confidence decrease with increasing uncertainty levels (Quiroga-Martinez et al., 2020a, 2020b, 2019). These effects can be interpreted in the light of predictive processing theories (Clark, 2013; Friston, 2009; Ross and Hansen, 2016; Vuust et al., 2018), suggesting that the salience of unexpected events and the neural responses they elicit are modulated by the certainty or precision of predictions. Thus, in uncertain auditory environments, where predictions are less precise, prediction error responses are attenuated (Garrido et al., 2013; Hsu et al., 2015; Sohoglu and Chait, 2016; Southwell and Chait, 2018).

The influence of uncertainty on predictive processing is particularly interesting in the case of atonal music. While Western tonal music possesses a tonal hierarchy in which certain tones are perceived as more stable and expected than others (Krumhansl and Cuddy, 2010), atonal compositions strive to avoid such a hierarchical structure and are based on a musical system in which tones tend to occur with equal probabilities. In other words, atonal music has a high degree of tonal uncertainty, evokes weaker expectancies in the listener, and is difficult to predict (Krumhansl et al., 1987; Mencke et al., 2019; Ockelford and Sergeant, 2012; Vuvan et al., 2014). Therefore, atonal music is a promising tool for studying how predictive models arise in high-uncertainty contexts.

In the present study, we employed a behavioral task and a multi-feature MMN paradigm targeting the influence of tonal uncertainty on prediction error responses. Specifically, we addressed the question of whether tonal uncertainty modulates the local auditory deviance processing indexed by the MMN and whether such modulation is reflected in behavioral deviance detection. Our stimuli were matched in a way that ruled out other sources of uncertainty such as the number of utilized pitches (pitch alphabet) and the degree of pitch pattern repetition. This allowed us to reliably isolate contextual tonal uncertainty effects. We hypothesized that for pitch deviants in the atonal condition, not only the MMN but also accuracy and confidence ratings in the behavioral task would be reduced in the atonal condition due to the high uncertainty of such stimuli. Surprisingly, however, our findings revealed a more complex picture in which low-level and higher order expectations were clearly differentiated.

## 2. Results

### 2.1 Presence of the MMNm for Different Features and Conditions

An MMNm was reliably evoked in both conditions and for all features (all p < .001 except for the pitch MMNm in the atonal condition p = .011; see Figure 1). In the right hemisphere, the magnetic gradient was positive at anterior sites and negative at posterior channels, whereas the left hemisphere showed the opposite pattern. This is a typical scalp topography for the MMNm (Quiroga-Martinez et al., 2019; Bonetti et al, 2017) (see Figure 2).

**Figure 1.**
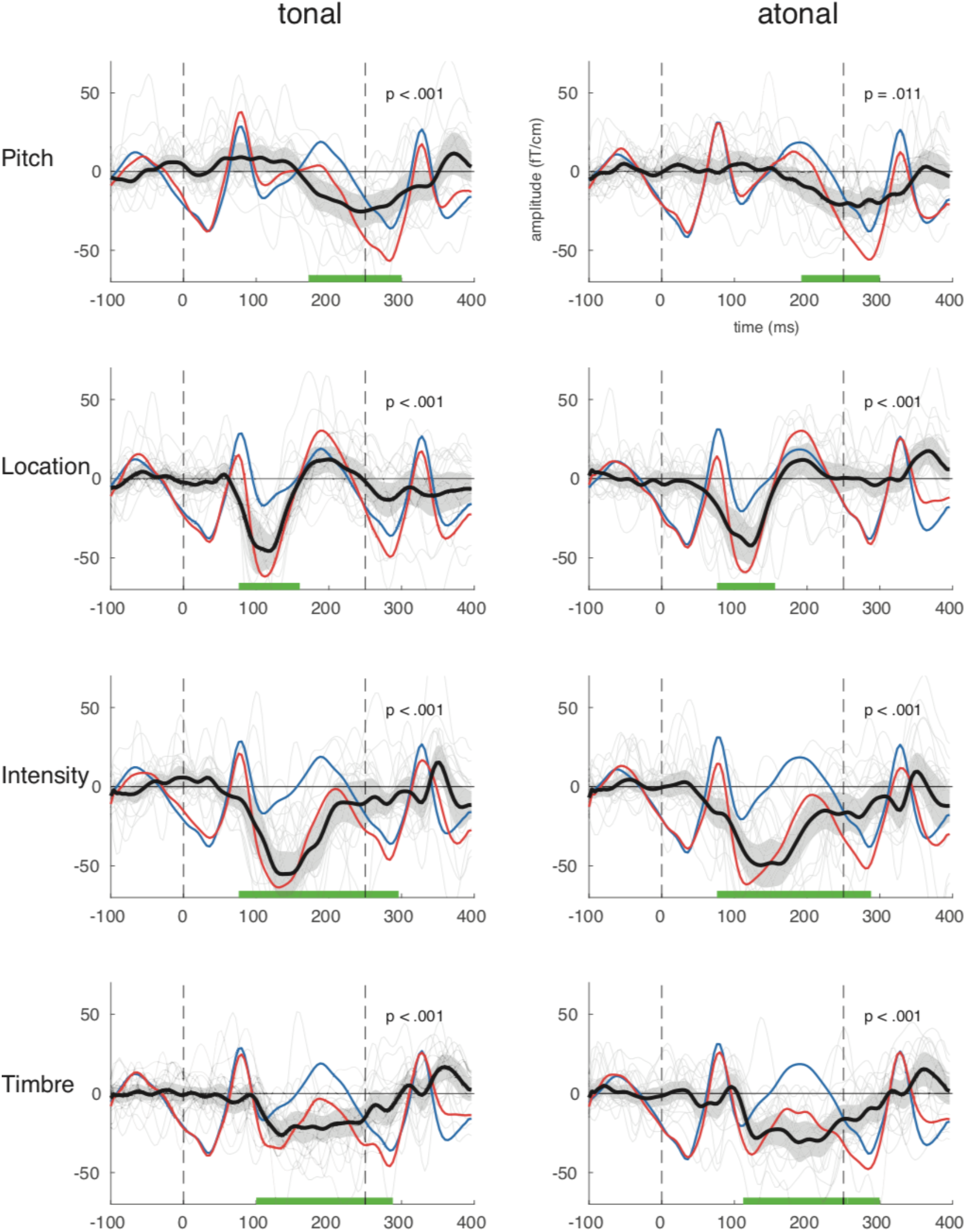
Average evoked responses to standard tones (blue), deviant tones (red) and difference waves (MMN; black) in the two conditions for all features. Green bars represent the time windows during which the differences between standard and deviant were significant. The dashed horizontal lines represent sound onsets. Shaded areas represent 95% confidence intervals. Grey lines in the background depict MMN time course for each participant. The displayed activity corresponds to the average of the following representative auditory channels on the right hemisphere: ‘MRT14’, ‘MRT15’, ‘MRT16’, ‘MRP55’, ‘MRP56’, ‘MRP57’.

**Figure 2.**
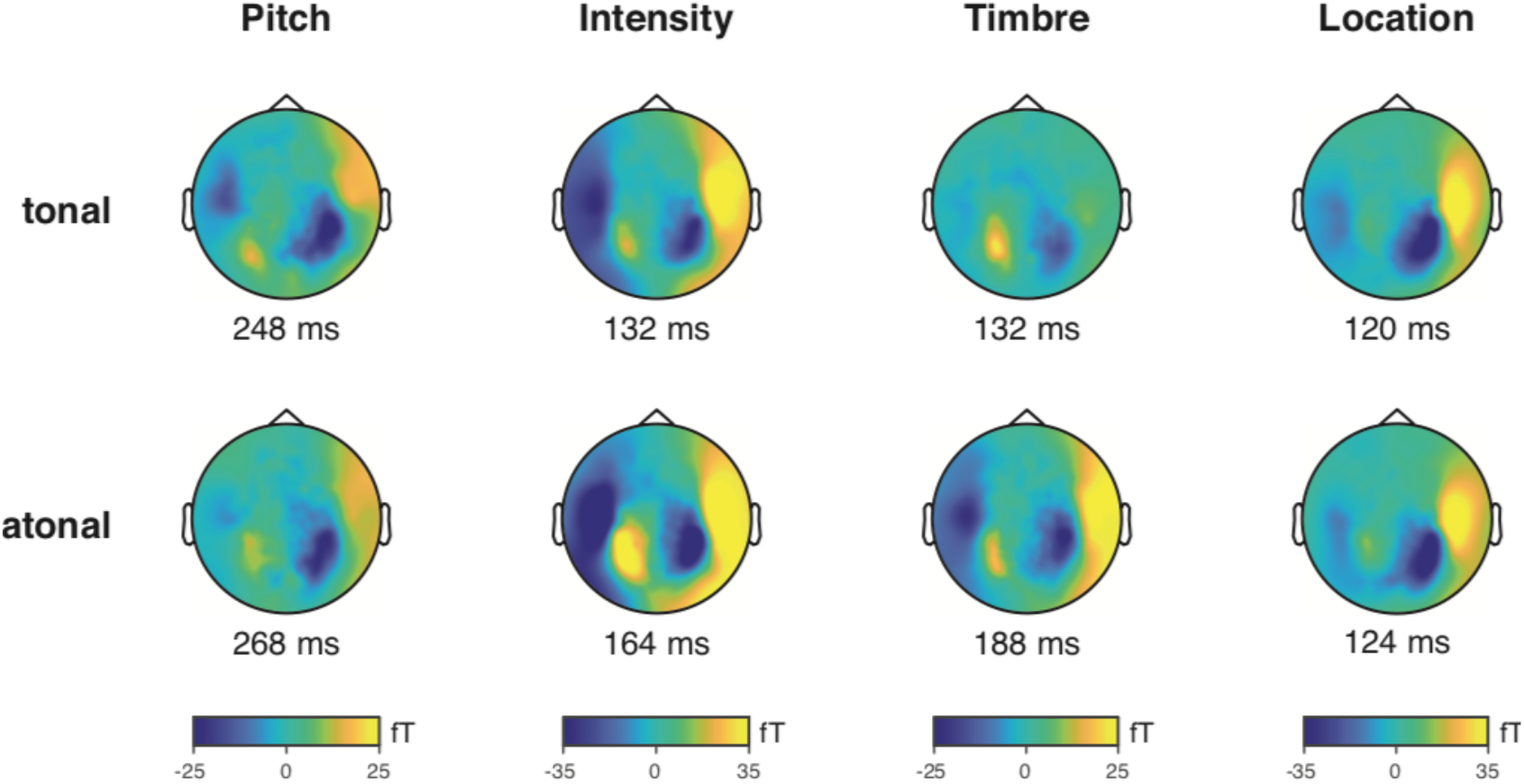
Topography of the MMNm (difference between standards and deviants). The displayed activity corresponds to an average of ±25 ms around the peak (latency shown below each plot).

### 2.2 Effects of Tonal Uncertainty

Contrary to our hypothesis, there was no significant change in MMNm amplitude between conditions for any of the four deviants (see Figure 3). This was confirmed in mixed-effects modeling of mean amplitudes, which revealed no significant effects of tonal uncertainty (see Table 1, *m1*). However, we found effects of the factors deviant feature (*m2*) and hemisphere (*m3*), whose inclusion significantly improved model performance. A significant deviant-by-hemisphere interaction (see Table 1, *m6*) was also present. Post-hoc pairwise contrasts between conditions for mean amplitudes yielded no significant differences for any feature or hemisphere (see Table 2).

**Figure 3.**
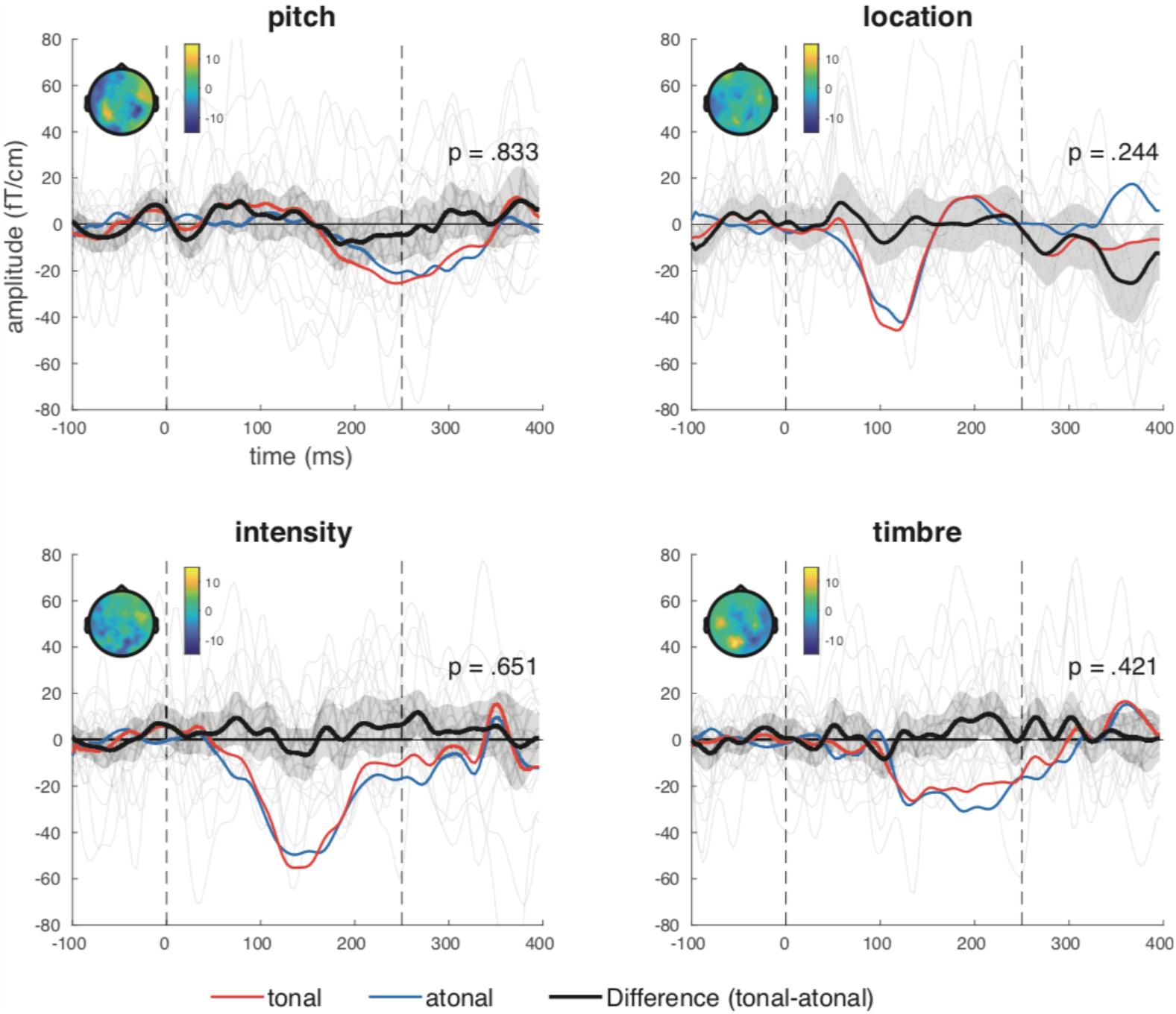
MMNm responses in the tonal (red) and the atonal conditions (blue), and the difference between them (black). Shaded areas depict 95% confidence intervals. Gray traces in the background depict the difference between conditions for each participant. The displayed activity corresponds to the average of the following representative auditory channels sensors on the right hemisphere: ‘MRT14’, ‘MRT15’, ‘MRT16’, ‘MRP55’, ‘MRP56’, ‘MRP57’.

**Table 1.** Likelihood ratio tests for mixed-effects models of mean amplitudes and peak latencies

| model | added term | null | Mean Amplitudes |  |  | Peak Latencies |  |  |
| --- | --- | --- | --- | --- | --- | --- | --- | --- |
| | | | AIC | $\chi^2$ | p | AIC | $\chi^2$ | p |
| m0 | intercept | NA | 2699.30 | NA | NA | 3315.94 | NA | NA |
| m1 | condition | m0 | 2701.09 | 0.21 | 0.65 | 3316.91 | 1.03 | 0.31 |
| m2 | deviant | m1 | 2577.08 | 130.02 | **<.001\*** | 3154.61 | 168.33 | **<.001\*** |
| m3 | hemisphere | m2 | 2574.25 | 4.82 | **0.03\*** | 3154.95 | 1.65 | 0.20 |
| m4 | cond:dev | m3 | 2575.66 | 4.60 | 0.20 | 3157.42 | 3.54 | 0.32 |
| m5 | cond:hem | m4 | 2577.48 | 0.18 | 0.67 | 3159.40 | 0.02 | 0.89 |
| m6 | dev:hem | m5 | 2575.53 | 7.94 | **0.05\*** | 3155.19 | 10.21 | **0.02\*** |
| m7 | cond:dev:hem | m6 | 2581.19 | 0.35 | 0.95 | 3160.17 | 1.02 | 0.80 |

*Note.* Models were built stepwise by adding one factor at a time and compared with the previous “null” model. Colons between factors indicate interactions. Moreover, Akaike information criteria (AIC), chi-square statistics ( $\chi^2$ ), and *p*-values (*p*) are reported. Significant likelihood ratio tests are marked with an asterisk.

**Table 2.** Pairwise Contrasts of Mean Amplitudes Between Tonal and Atonal Conditions by Deviant Feature and Hemisphere.

| contrast | deviant | hemisphere | estimate | CI 2.5% | CI 97.5% | t | p | d |
| --- | --- | --- | --- | --- | --- | --- | --- | --- |
| tonal - atonal | pitch | left | -4.62 | -14.53 | 5.28 | -0.92 | 0.36 | -0.31 |
| tonal - atonal | location | left | 1.31 | -8.59 | 11.22 | 0.26 | 0.79 | 0.09 |
| tonal - atonal | intensity | left | -3.59 | -13.50 | 6.32 | -0.71 | 0.48 | -0.24 |
| tonal - atonal | timbre | left | 5.82 | -4.09 | 15.72 | 1.16 | 0.25 | 0.39 |
| tonal - atonal | pitch | right | -5.67 | -15.58 | 4.23 | -1.13 | 0.26 | -0.38 |
| tonal - atonal | location | right | -3.21 | -13.12 | 6.69 | -0.64 | 0.52 | -0.21 |
| tonal - atonal | intensity | right | -2.38 | -12.28 | 7.53 | -0.47 | 0.64 | -0.16 |
| tonal - atonal | timbre | right | 4.21 | -5.70 | 14.12 | 0.84 | 0.40 | 0.28 |

*Note.* Confidence intervals (CI), *t*-values (t), *p*-values (*p*) and effect sizes (d) are reported.

We also conducted mixed-effects modeling of peak latencies. The results revealed no significant effect of tonal uncertainty (see Table 1, *m1*) but once more significant effects of deviant feature (*m2*) and hemisphere (*m3*) were observed. As for mean amplitudes, a significant deviant-by-hemisphere interaction (*m6*) was also present. Post-hoc pairwise contrasts between conditions for peak latencies (see Table 3) did not reveal any significant effects.

**Table 3.** Pairwise contrasts of peak latencies between tonal and atonal conditions by deviant feature and hemisphere.

| contrast | deviant | hemisphere | estimate | CI 2.5% | CI 97.5% | t | p | d |
| --- | --- | --- | --- | --- | --- | --- | --- | --- |
| tonal - atonal | pitch | left | 9.05 | -17.70 | 35.81 | 0.67 | 0.51 | 0.22 |
| tonal - atonal | location | left | -5.05 | -31.81 | 21.71 | -0.37 | 0.71 | -0.12 |
| tonal - atonal | intensity | left | -11.79 | -38.54 | 14.96 | -0.87 | 0.39 | -0.29 |
| tonal - atonal | timbre | left | -20.84 | -47.60 | 5.91 | -1.53 | 0.13 | -0.51 |
| tonal - atonal | pitch | right | 7.37 | -19.38 | 34.12 | 0.54 | 0.59 | 0.18 |
| tonal - atonal | location | right | -17.89 | -44.65 | 8.86 | -1.32 | 0.19 | -0.44 |
| tonal - atonal | intensity | right | -2.53 | -29.28 | 24.23 | -0.19 | 0.85 | -0.06 |
| tonal - atonal | timbre | right | -10.32 | -37.07 | 16.44 | -0.76 | 0.45 | -0.25 |

*Note.* Confidence intervals (CI), *t*-values (t), *p*-values (*p*) and effect sizes (d) are reported.

### 2.3 Feature and Hemisphere Specific Differences

Post-hoc, Bonferroni-corrected pairwise contrasts for mean amplitudes between features (see Table 4) revealed that in both hemispheres all feature-specific differences were significant (see Table 1, *m2*). The pitch deviant evoked smallest MMN amplitude in contrast to all other features but especially in contrast to the intensity deviant. Location was weaker than intensity and timbre was weaker than both location and intensity. With regard to peak latencies a similar pattern of results emerged. The pitch deviant occurred later than all other deviants and timbre occurred later than location and intensity deviants. However, the location-intensity contrast for peak latencies revealed that the location deviant was evoked earlier (see also Figure 4).

**Table 4.** Pairwise contrasts of mean amplitudes and peak latencies between features by hemisphere.

| contrast | Mean amplitudes |  |  |  |  |  | Peak Latencies |  |  |  |  |  |
| --- | --- | --- | --- | --- | --- | --- | --- | --- | --- | --- | --- | --- |
|  | estimate | CI 2.5% | CI 97.5% | t | p | d | estimate | CI 2.5% | CI 97.5% | t | p | d |
| pitch - location | 15.95 | 9.26 | 22.63 | 6.34 | <.001* | 1.06 | 95.63 | 77.58 | 113.68 | 14.07 | <.001* | 2.34 |
| pitch - intensity | 31.08 | 24.40 | 37.77 | 12.35 | <.001* | 2.06 | 76.95 | 58.89 | 95.00 | 11.32 | <.001* | 1.89 |
| pitch - timbre | 08.08 | 1.39 | 14.76 | 3.21 | **0.01\*** | 0.53 | 43.26 | 25.21 | 61.32 | 6.36 | <.001* | 1.06 |
| location - intensity | 15.14 | 8.45 | 21.82 | 6.01 | <.001* | 1.00 | -18.68 | -36.74 | -0.63 | -2.75 | **0.04\*** | -0.46 |
| location - timbre | -7.87 | -14.55 | -1.19 | -3.13 | **0.01\*** | -0.52 | -52.37 | -70.42 | -34.32 | -7.70 | <.001* | -1.28 |
| intensity - timbre | -23.01 | -29.69 | -16.32 | -9.14 | <.001* | -1.52 | -33.68 | -51.74 | -15.63 | -4.96 | <.001* | -0.83 |

*Note.* Confidence intervals (CI), *t*-values (t), *p*-values (*p*) and effect sizes (d) are reported. Significant contrasts are marked with an asterisk.

**Figure 4.**
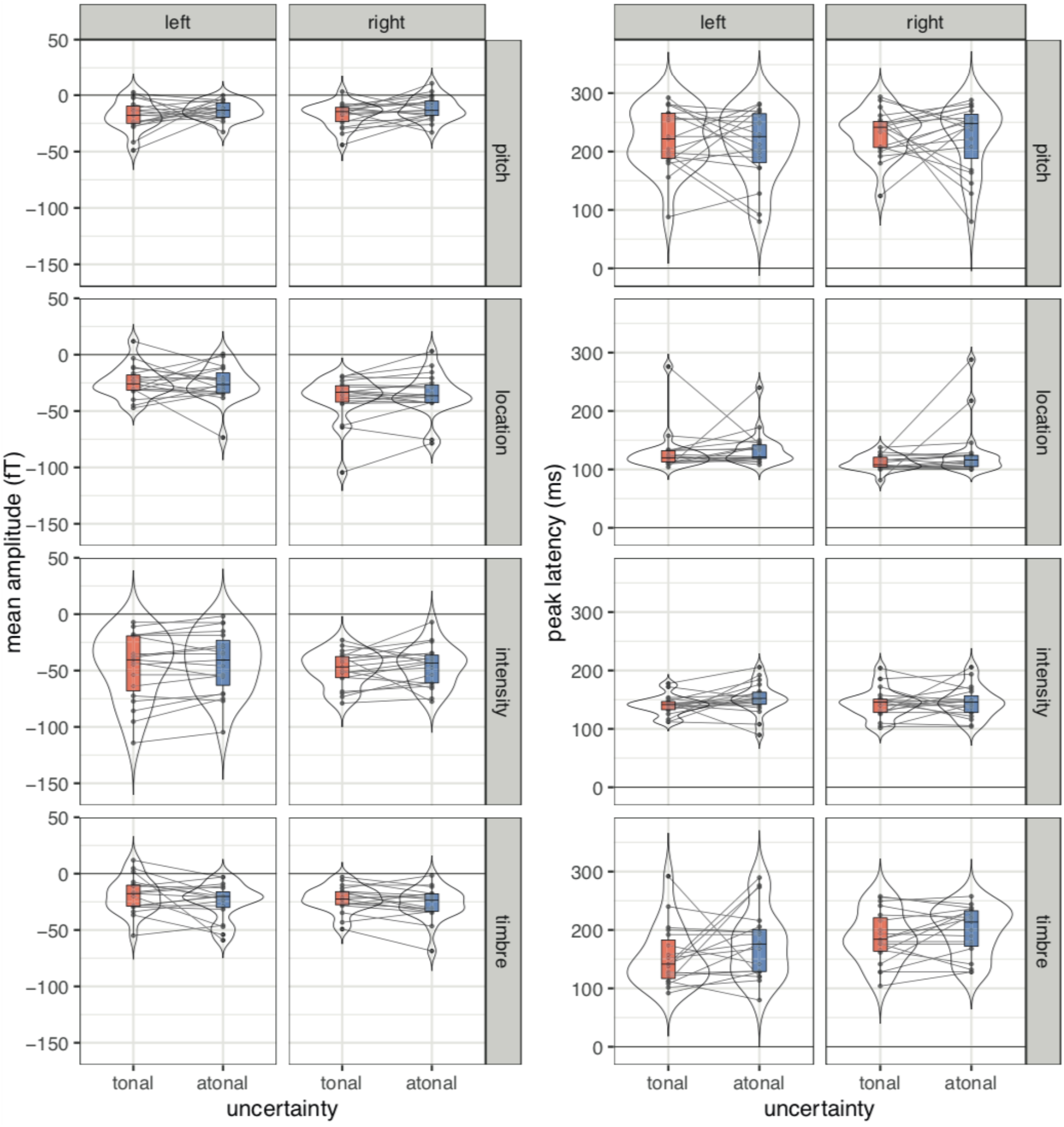
Mean MMN amplitudes (left) and peak latencies (right) as a function of uncertainty in both hemispheres. Boxplots display median and interquartile ranges. Beans depict the estimated densities. Lines connect measurements for individual participants. Displayed data were obtained from selected auditory channels on each hemisphere (see main text for details).

The hemisphere differences (Table 5) show that in the case of mean amplitudes the effect of hemisphere (*m3*) was due to the location deviant being significantly stronger in the right hemisphere, and in the case of latencies it was due to the timbre deviant being significantly stronger in the left hemisphere.

**Table 5.** Pairwise contrasts of mean amplitudes and peak latencies between hemispheres for features.

| contrast | feature | Mean amplitudes |  |  |  |  |  | Peak latencies |  |  |  |  |  |
| --- | --- | --- | --- | --- | --- | --- | --- | --- | --- | --- | --- | --- | --- |
|  |  | estimate | CI 2.5% | CI 97.5% | t | p | d | estimate | CI 2.5% | CI 97.5% | t | p | d |
| left - right | pitch | -1.04 | -8.05 | 5.96 | -0.29 | 0.77 | -0.07 | -10.74 | -29.65 | 8.18 | -1.12 | 0.26 | -0.71 |
| left - right | location | 12.09 | 05.08 | 19.09 | 3.40 | <.001* | 0.80 | 12.32 | -6.60 | 31.23 | 1.28 | 0.20 | 0.82 |
| left - right | intensity | 1.98 | -5.03 | 8.98 | 0.56 | 0.58 | 0.13 | 1.89 | -17.02 | 20.81 | 0.20 | 0.84 | 0.13 |
| left - right | timbre | 2.61 | -4.39 | 9.62 | 0.73 | 0.46 | 0.17 | -28.21 | -47.13 | -9.29 | -2.93 | <.001* | -1.87 |

*Note.* Confidence intervals (CI), *t*-values (t), *p*-values (*p*) and effect sizes (d) are reported. Significant contrasts are marked with an asterisk.

### 2.4 Source Reconstruction

For all features, we found significant differences between standard and deviants (all p < .001) with, in most cases, clear peaks in the surroundings of the auditory cortex. Likely due to the lower signal-to-noise ratio of pitch and timbre MMNm responses, the auditory peaks of these features were closer in amplitude to occipital differences, which likely reflect visual artifacts related to watching the movie that did not completely cancel out after averaging and subtraction. No clear frontal peaks were observed for any features. Peak coordinates for left and right hemispheres are reported in supplementary material 3.

### 2.5 Behavioral Experiment

Sensitivity (*d*’) scores were significantly lower in the atonal than the tonal condition (t(38) = 5.15, p < .001, M = -0.65, CI (95%) = [-0.9,-0.39]), whereas there were no significant differences in *c-*scores as a measure of bias (t(38) = 0.69, p = .49, M = -0.07, CI (95%) = [-0.26, 0.13]) (see Figure 6*A* and 6*B*). The ordinal regression model revealed that confidence ratings were lower for atonal as compared with tonal deviant melodies (OR = 0.54, p < .001, CI (95%) = [0.43, 0.69]), and that regular tonal melodies were not rated significantly different from deviant tonal melodies (OR = 0.87, p = .266, CI (95%) = [0.69, 1.11]; see Figure 6*C, B*). There was no significant deviance-by-condition interaction either (OR = 0.78, p = .136, CI (95%) = [0.57, 1.08]). However, in the extended analyses with all the participants included, this interaction became significant (OR = 0.76, p = .048, CI (95%) = [0.58, 0.997]) (see supplementary material 4 for a full report). This can already be seen in Figure 6*C*, which shows that the difference in ratings between atonal and tonal melodies was even more pronounced in the regular as compared with the deviant melodies.

**Figure 5.**
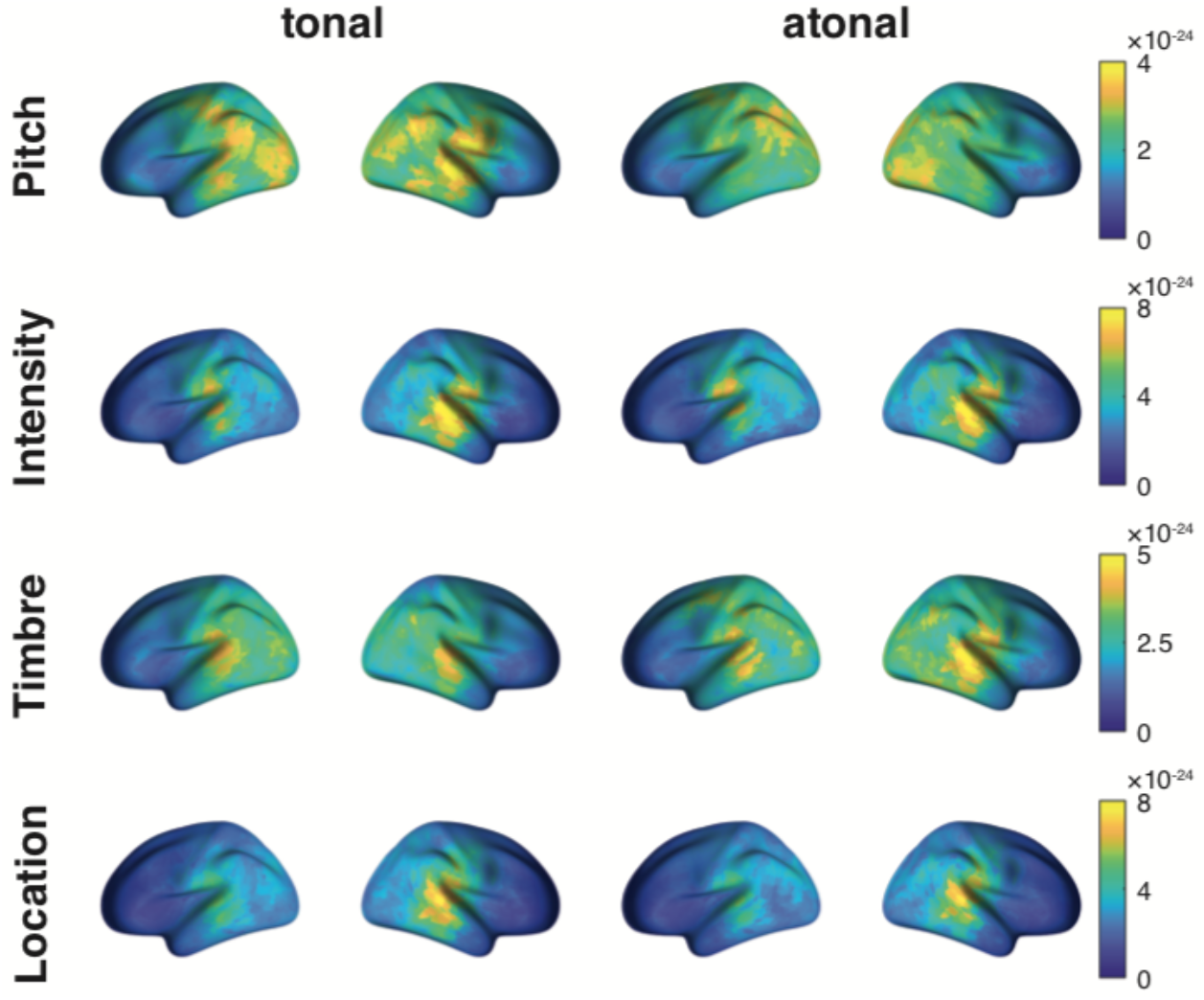
MMNm source power for each deviant feature and condition. Displayed activity corresponds to the grand-average source estimates of the difference between standards and deviants ±25 ms around each participant’s sensor-level MMNm peak. Peak coordinates for left and right hemispheres are reported in supplementary material 3.

**Figure 6.**
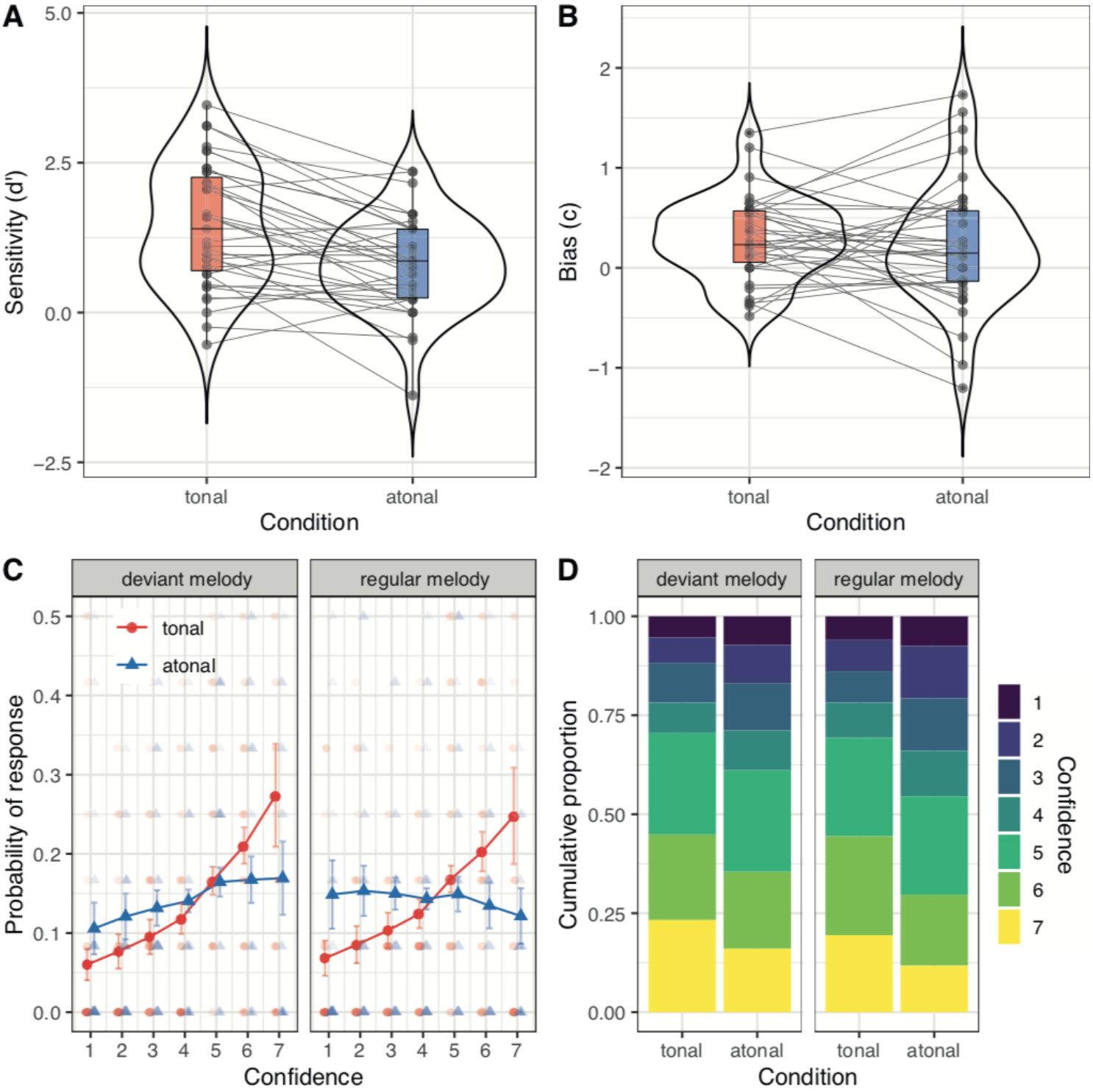
Behavioral measures of pitch deviance detection. **A**, Sensitivity scores. **B**, Bias estimated as criterion (*c*) scores. ***C***, Estimated probability that participants will choose a particular confidence rating (from 1 to 7), according to the two conditions. Note how lower ratings increase, and higher ratings decrease, in atonal as compared with tonal melodies, and how this effect is stronger for regular than for deviant melodies. Small dots represent individual data points (values above 0.5 not shown). ***D***, Cumulative proportion of individual responses for each confidence rating category. See supplementary material 4 for a full report and supplementary material 5 for a report with the full sample of participants included.

Based on these results, we further explored whether participants’ accuracy followed a similar pattern by estimating a mixed-effects logistic regression model with condition, deviance, and their interaction as predictors. We found significantly lower accuracy for atonal as compared with tonal deviant melodies (OR = 0.73, p = .023, CI (95%) = [0.56, 0.96], significantly higher accuracy for regular as compared with deviant tonal melodies OR = 2.72, p < .001, CI (95%) = [2, 3.71]), and a non-significant deviance-by-condition interaction (OR = 0.69, p = .085, CI (95%) = [0.46, 1.05]). As with confidence ratings, this interaction became significant once the full sample of participants was analyzed (OR = 0.71, p = .049, CI (95%) = [0.51, 0.999]) (see supplementary material 5). Overall, these results indicate that participants are less accurate and become less confident at detecting deviant sounds in atonal as compared with tonal melodies. Moreover, they also suggest that accuracy and confidence further decrease in the presence of atonal melodies that do not contain a deviant, thus highlighting the behavioral difficulty of distinguishing regular from deviant sounds in high-uncertainty contexts.

## 3. Discussion

Our data showed that the MMNm for intensity, location, pitch, and timbre was relatively unaffected by tonal uncertainty, and that MMNm responses were robustly evoked in both tonal and atonal conditions. Strikingly, however, the behavioral results indicated that detection of pitch deviants and subjective confidence were reduced in the atonal context. Our results point to a differentiation between early latency neural and slower behaviorally measured signatures of music-predictive processing in contexts of high uncertainty: despite the listeners’ reduced awareness as indicated by the behavioral findings, sensory deviants were reliably detected in low-level, pre-attentive processing stages.

### 3.1 Low-level and High-Level Predictive Processing Stages

In agreement with Quiroga-Martinez et al. (2019), we found MMNm responses for all deviant features in the tonal melodies. However, in contrast to that study, we did not find an effect of contextual tonal uncertainty on the amplitude or latency of the MMNm. Therefore, our results suggest that tonal uncertainty does not influence the degree to which deviants are processed at the low-level stages reflected by the MMN. However, the behavioral experiment, which engaged processes requiring conscious awareness (Dehaene and Changeux, 2011), showed that deviance-detection performance and confidence ratings were significantly lower for atonal as compared with tonal melodies.

The MMN is not necessarily modulated by the conscious awareness of deviants (Garrido et al., 2009; Näätänen et al., 2007). Rather, it is associated with pre-attentive sensory processing and the extraction of low-level regularities (Koelsch, 2012; Näätänen, 1992) and is therefore responsive to events that are locally salient (Bekinschtein et al., 2009; King et al., 2014; Peretz et al., 2009; Wacongne et al., 2012). In our study, we violated regularities of deviant features that were mainly sensory and therefore equally salient in both conditions. Especially location (interaural differences) and intensity deviants are known to be processed relatively early in the auditory pathway, namely in the superior olivary complex (Schnupp et al., 2011), which may explain the large amplitude of these MMNm responses. As argued by Brattico and colleagues (2006), these deviant features likely induce the MMN due to their extreme rareness, which was matched across our conditions (see also Saarinen et al., 1992). Moreover, a recent study suggests that a physical, i.e., sensory MMN is so strong that it overrules a “statistical MMN” which reflects larger scale manipulations (Tsogli et al., 2019). Our finding parallel those from other studies in which early mismatch responses were relatively intact in subjects with congenital amusia, a disorder of musical listening (Omigie et al., 2013; Peretz et al., 2009, 2005; Quiroga-Martinez et al., 2021). In this population, only later P600 responses indicated atypical processing of pitch deviants (Moreau et al., 2013; Peretz et al., 2009; Zendel et al., 2015), while the MMN was reliably evoked.

Since the MMNm is not a direct neural correlate of behavioral deviance detection, it is likely that other neural responses reflect tonal uncertainty effects on behavior. One candidate is the P300, which is associated with conscious novelty detection (King et al., 2014, 2013) and which occurs significantly later than the MMN. It responds to violations of global auditory expectations that are processed at higher levels (Bekinschtein et al., 2009; Wacongne et al., 2012). The P300 comprises two subcomponents, the P3a and P3b (Polich, 2007; Polich and Criado, 2006), which are said to be particularly modulated by the uncertainty of the stimulus (Kopp and Lange, 2013; Sutton et al., 1965).

One could speculate that the lower detection of deviants and confidence ratings in the atonal condition would be reflected in an attenuation of the P3b. However, it is possible that our manipulation would only induce such an attenuation when participants are consciously aware of the stimuli (King et al., 2014, 2013). In this regard, it is important to note that the stimuli in the MEG experiment were task-irrelevant and unattended. Moreover, due to the tone duration of 250 ms we could not precisely measure such late evoked components since they would overlap with the evoked response of the next tone. Future studies should address the role of attentive versus non-attentive predictions and prediction errors since this may potentially reveal how the processing of tonal hierarchies is shaped by attention.

### 3.2 Effects of Tonal Uncertainty on the MMN

We employed stimuli in which the two sources of uncertainty manipulated in previous research were controlled in order to isolate tonal uncertainty effects. First, we matched our two conditions in their pitch-alphabet, which contrasts with the stimuli of Quiroga-Martinez and colleagues (2019), whereby the low-entropy condition was restricted to an alphabet of three pitches, while the high-entropy condition had a much larger alphabet, similar to ours. Thus, our results suggest that pitch-alphabet size plays an important role in modulating MMN amplitude. The reduction of neural responses with increasing alphabet sizes has previously been found for non-musical stimuli (Barascud et al., 2016; Herrmann et al., 2014). Similarly, it has been shown that neuronal populations modulate their excitability depending on the variance of sensory information, effectively adapting to the statistics of the stimuli (Wark et al., 2007; Weber et al., 2019). Therefore, the dampening of neural responses to deviant sounds by larger alphabet sizes might reflect the fact that more auditory information needs to be kept in sensory memory, resulting in competition for neural resources and mutual inhibition. This can be seen as a basic form of precision-weighting acting on low-level sensory representations instead of the higher-order statistical regularities underlying tonal uncertainty.

Another related property that we controlled for is the presence of exact pattern repetitions in the stimuli. The few pitches in the low-entropy condition in Quiroga-Martinez and colleagues (2019) were organized into a short melodic pattern that was repeated throughout the stimulation. This likely established a strong predictive model, not only with regard to single pitches, but also with regard to the pattern itself. This short and highly repetitive type of stimulation has been the common way of assessing MMNs in musical contexts (Fujioka et al., 2004; Tervaniemi et al., 2001; Zuijen et al., 2004). In contrast, the present study used melodies with a constantly changing melodic contour in both the high- and low-uncertainty conditions. Our results suggest that, after controlling for factors such as pitch alphabet and the presence of exact pattern repetitions, which operate on lower processing levels, there is little or no effect of tonal uncertainty on the MMNm.

### 3.3 Feature- and Hemisphere-Specific Effects

In our study, pitch deviants resulted in the smallest mean amplitudes and the longest latencies (see Table 4), in agreement with previous musical MMN paradigms for the same deviant magnitudes and subject groups (Vuust et al., 2016). Moreover, similar to the pitch deviant, the timbre deviant showed a relatively small amplitude (see Figures 3 and 4), which also corroborates previous findings (Kliuchko et al., 2019). In fact, in musical multi-feature paradigms, deviants of different features tend to have different levels of salience (Vuust et al., 2016). This, however, does not interfere with the tonal uncertainty manipulations made here. Lastly, in pairwise comparisons we observed a right-lateralization of the location deviant with regard to mean amplitude (see Table 5 and Figure 3). This effect was likely driven by the leftward location bias which resulted in the MMNm being lateralized to the opposite hemisphere (Richter et al., 2009; Sonnadara et al., 2006).

### 3.4 Limitations and Future Directions

First, since our study only tested non-musicians, an interesting question is how musically trained individuals would respond to the two conditions. As multiple studies have shown, the MMN to deviating sounds in an auditory sequence is significantly enhanced in musicians as compared with non-musicians (Tervaniemi et al., 2014; Vuust et al., 2012, 2011) and even discriminates between different types of musicians (Vuust et al., 2012). Expertise effects on the P3b in response to unexpected stimuli have been observed (Przysinda et al., 2017), whereas a recent study did not find an interaction between expertise and entropy (Quiroga-Martinez et al., 2019). However, testing musicians with our more carefully controlled stimuli may reveal whether expertise-related interactions exist in a more ecological, high-uncertainty musical context.

Moreover, although our stimuli had a high level of ecological validity in contrast to previous musical multi-feature MMN studies, their implications for naturalistic listening situations are restricted since they are still rather artificial with respect to the way they were generated and presented. In fact, atonal music occurs in numerous forms and varieties (Mencke et al., 2019) and the lack of tonal hierarchies is often “compensated for” with the presence of other forms of hierarchies based on timbre or rhythmical aspects, for instance (Bharucha, 1984; Brattico et al., 2017).

While previous research focused on paradigms often using stimuli composed according to Western tonal rules, stimulation paradigms have only recently become more complex and ecological (Haumann et al., 2021; Poikonen et al., 2016; Quiroga-Martinez et al., 2019). With respect to studies in music cognition, our study highlights the relevance of including a larger variety of musical styles or musical vocabulary when investigating expectancy processes. More generally, our paradigm and results should contribute to a better understanding of how to study complex stimuli in a controlled way.

Finally, our study provides new research avenues for the investigation of mechanisms that underly engagement with uncertainty. In daily life, human individuals are often exposed to novel environments in which predictions are imprecise and in which learning is required. It is therefore of crucial importance to better understand how predictions are being generated in such environments and what mechanisms lead to an increase in the strength of predictions. Studying atonal music or similar sound contexts that comprise a low degree of predictability could contribute to a better understanding of such predictive processes under high uncertainty circumstances.

### 3.5 Conclusion

This study gives insights into the consequences that the lack of a regular and hierarchically structured input has on neural and cognitive processing. Our behavioral results suggest that the generation of predictive models is strongly impeded on higher processing levels. In contrast, the lower levels indexed by the MMN are not affected by a lack of tonal hierarchies. This suggests that (at least) two distinctive types of expectations may be at work during music listening. Thus, presenting stimuli that entail a high predictive uncertainty can help to establish a more fine-grained picture of predictive neuronal processing, which may contribute to a deeper understanding of how different types of predictive models evolve in the brain.

## 4. Materials and Methods

### 4.1 Participants (MEG experiment)

Twenty healthy right-handed subjects (mean age = 31.6; *SD* = 5.7; female = 13) were recruited via an online recruitment platform of the Max Planck Institute for Empirical Aesthetics (Frankfurt/Main, Germany). Additional questions in the invitation process were asked to confirm that participants fulfilled the criteria for the required levels of musical training (1. No instrumental training before the age of ten; 2. No instrumental training longer than three years). This was controlled by assessing the participants’ musical backgrounds with the general factor of the Goldsmiths Musical Sophistication Index (Schaal et al., 2014) in the second session. The mean score was 50.2 (*SD* = 17.4; *Mdn* = 48.5), which equals the lowest 7% of the norm scale (minimum = 32; maximum = 126). All participants gave written informed consent in advance of the study and received monetary compensation at the end. The data from all participants were included in the analyses. The study was conducted according to the Declaration of Helsinki and was approved by the local ethics committee of the University Hospital Frankfurt (reference number 415/17). Individual MRI scans were recorded for all participants except for one who did not fulfill the MRI inclusion criteria as they only later reported having worked in the metal industry. Another participant was excluded from source localization analyses because their MRI was deficient.

### 4.2 Stimuli

#### 4.2.1 Generation of Melodies

The experiment consisted of tonal and atonal melodies that included four different kinds of deviants (see Figure 7). Tonal stimuli were adopted from previous studies (Quiroga-Martinez et al., 2020a, 2020b, 2019) and consisted of a set of six melodies composed according to the rules of Western tonal music. The melodies did not include exact internal repetitions of pitch patterns. Each individual melody consisted of 32 notes and had a duration of 8 seconds. Atonal stimuli were generated specifically for this study and similarly consisted of six different melodies adopted from twelve-tone rows from existing compositions (see Figure 8*C* for pitch class distributions in tonal and atonal melodies). Such rows were utilized in the context of the twelve-tone technique developed by Arnold Schönberg (Dibelius, 1998) at the beginning of the twentieth century with the underlying premise that all twelve notes within an octave would be treated equally. Specifically, each composition was based on a specific order of the twelve notes of the chromatic scale, in which an essential criterion was the avoidance of note repetitions, or, in other words, in which all twelve notes of the chromatic scale should appear once (Krumhansl, 1990). This basic sequence, called the prime form (Dienes and Longuet-Higgins, 2004), determines the specific order in which the tones appear in a composition. The prime form is modified in terms of three transformations that allow variations of the form while preserving its idiosyncratic character: the *retrograde* transformation reverses the order of the tones; the *inversion* inverts the pitch direction of the intervals; and the *retrograde inversion* inverts the temporal order of the inversion (Krumhansl and Cuddy, 2010). The first two formal transformations, namely the *retrograde* and the *inversion*, were built for each of the six prime forms. These three rows were then concatenated into one sequence which started with the prime form, was followed by the *retrograde* and the *inversion*. This procedure led to six atonal tone sequences of 36 notes.

**Figure 7.**
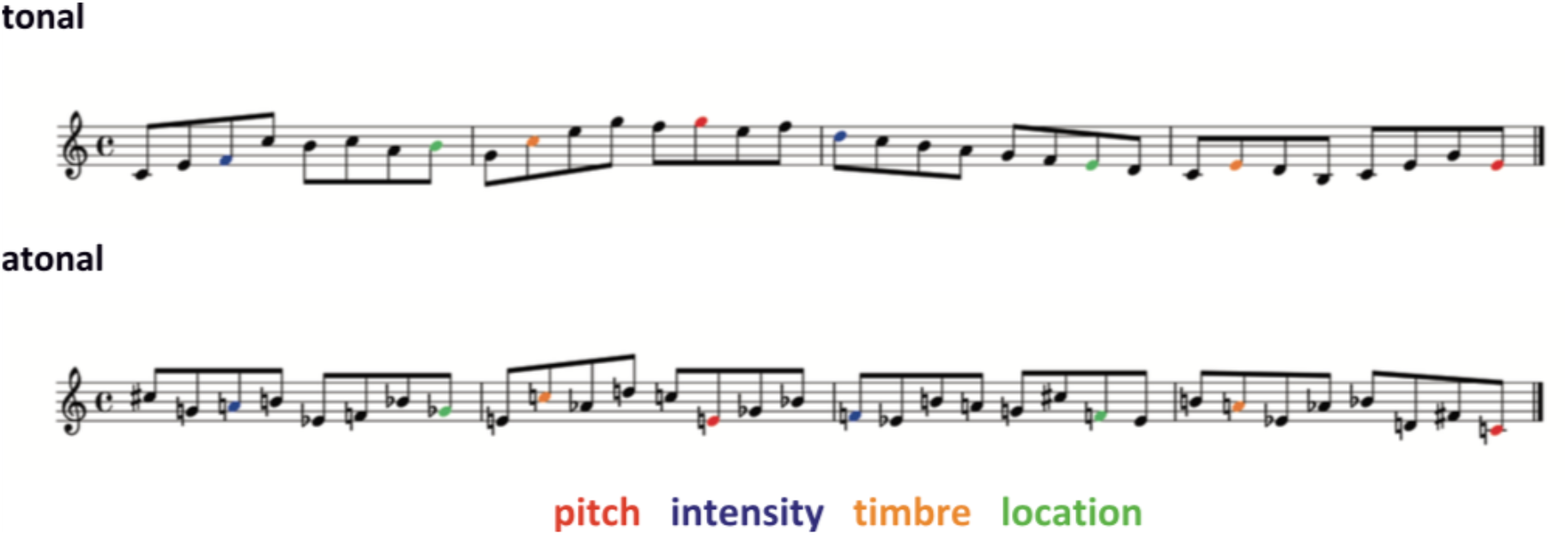
Examples of a 32-note melody from the tonal and the atonal conditions including the four different deviant types. The basis for the atonal melodies can be found in supplementary material 1.

**Figure 8.**
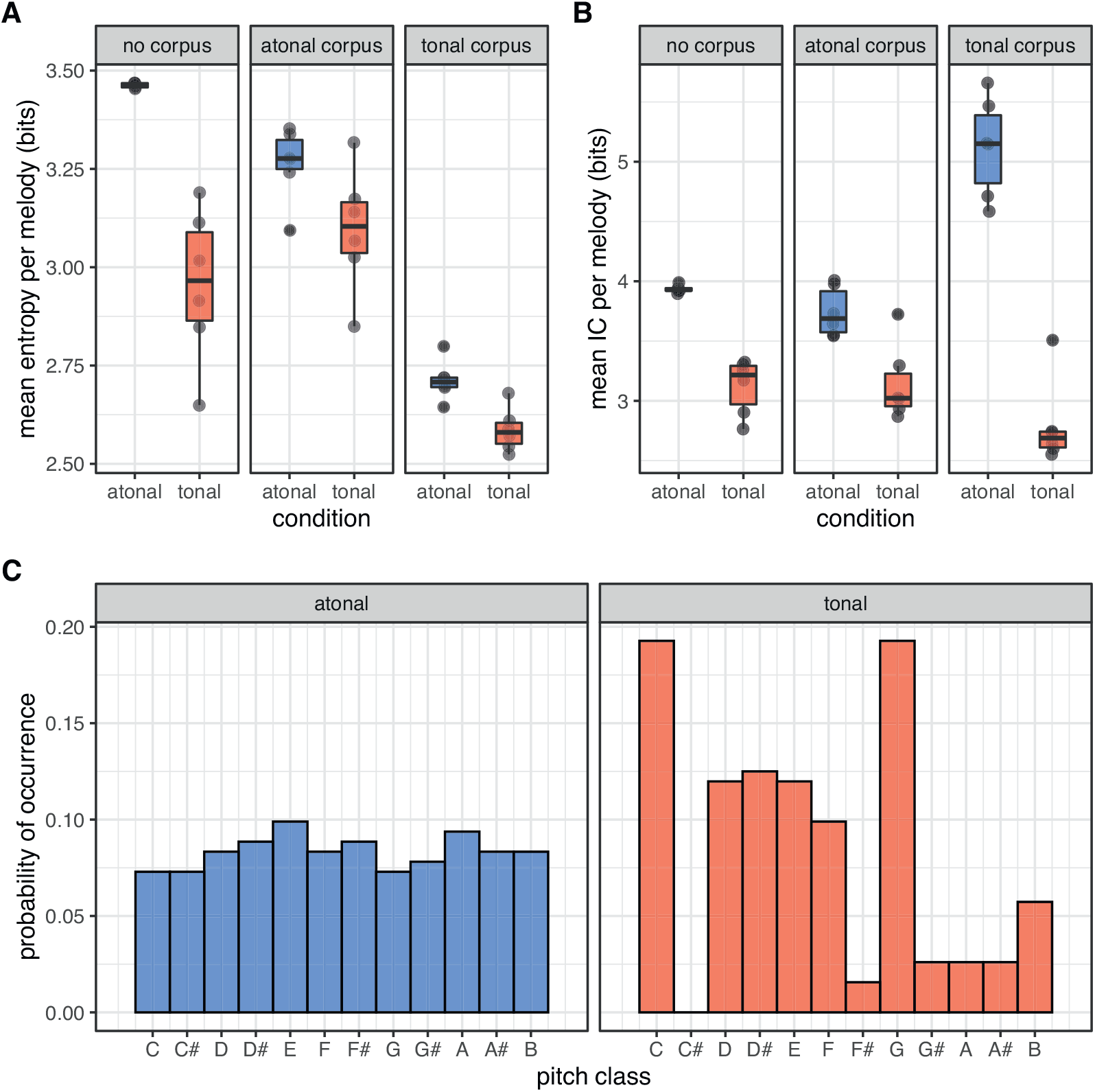
Computational measures of tonal uncertainty. ***A***, Mean entropy, and ***B***, Information content (IC) for each of the tonal and atonal melodies in our stimulus set, according to IDyOM models trained with either tonal, atonal, or no long-term corpora. Overall, atonal melodies have higher mean entropy and IC than tonal melodies. ***C***, Probability of occurrence of different pitch classes in tonal and atonal melodies in our stimulus set. Note how in tonal music the tonic (C) and the dominant (G) have a higher probability than the other notes, and how notes like F# and C# occur very rarely or not at all. By contrast, in atonal melodies pitch classes are distributed much more uniformly. For these analyses, tonal melodies we re transposed to a common key (C). For the exact pitch content included in both conditions see supplementary material 2.

In order to match the atonal and tonal melodies with regard to number of tones, each atonal sequence was reduced to 32 notes. This was done by following two criteria: 1) no consecutive pitch repetitions and 2) no consonant intervals may occur in the 32-tone melody. Thus, tones were deleted that corresponded to these criteria. Furthermore, we ensured that the tonal and the atonal melodies were matched in terms of pitch content (see supplementary material 2). The six resulting melodies can be found in supplementary material 1. The melodies in both conditions were pseudorandomly transposed from 0 to 5 semitones upwards. For tonal melodies, this corresponds to the keys F, F#, G, G#, A, and A#. For each of the keys both major and minor modes were used. After transpositions, the pitch range encompassed 31 semitones, from E3 (F_0_ ≈ 164 Hz) to B♭5 (F_0_ ≈ 932 Hz). In total, 72 melodies were played in each condition.

We used a grand piano sample from the Halion sampler in Cubase (Steinberg Media Technology, version 8) to generate the tone pool for stimulus presentation. Each note had a duration of 250 ms. Moreover, tones were normalized by peak amplitude and were subjected to a 3-ms-long fade-in and fade-out. The 250-ms tone duration shortened stimulation time while preventing the MMN from overlapping with the onset of the following tone. The individual melodies were not separated by gaps. Rather, they were continuously positioned in order to create the impression of real musical phrases. All the tones in the melodies were isochronous in order to ensure comparability both between conditions and with previous MMN paradigms.

#### 4.2.2 Introduction of Deviants

Four deviant types were generated using Audition (Adobe Systems Incorporated, version 8) and introduced in the atonal and tonal sequences. Pitch deviants were created by mistuning the pitch of the standard tone by raising it 50 cents. Intensity deviants were created by decreasing the volume of the tone by 12dB. Timbre deviants were generated by applying an old-time-radio filter available in Audition to the standard tone. Finally, location deviants were generated by applying a leftward bias (10 ms interaural time difference) to the standard tone.

The introduction of deviants followed a systematic pattern: Each melody was divided into eight groups of four notes each (see Figure 7), and a deviant occurred at random in any of the four notes of each group with equal probability. It was ensured that 1) none of the deviant features occurred consecutively, and 2) a deviant feature did not occur again before a whole iteration of the four features was played. The order of deviant features was pseudorandom. In total, 72 melodies were played in each condition, yielding 144 deviants per feature. The zeroth-order probability of each deviant type was 2/32 = 0.0625.

Each condition was presented in a separate block. At the beginning of each block, a melody without deviants was played to properly establish auditory regularities. Transpositions of the melodies were pseudorandomized in both conditions. Each block lasted approximately 11 minutes. The order in which the atonal and tonal blocks were presented was counterbalanced across participants.

### 4.2.3 Computational Measures of Tonal Uncertainty

To quantitatively assess tonal uncertainty in our stimulus set, we used Information Dynamics of Music (IDyOM; Pearce, 2005), a variable-order Markov model of musical expectations. IDyOM estimates the probability that a given tone will follow in a melody by counting the times that a given melodic pattern has been previously encountered in the current musical piece and/or in a training corpus. Thus, IDyOM reflects the statistical learning of regularities in musical pieces. The model has two output measures. The first is Shannon entropy:

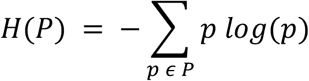

Here, *p* is the probability of a given tone continuation, given previous sound events. Note that entropy (*H*) is a direct measure of uncertainty, being largest when all continuations are equally probable and smallest when there is complete certainty that a particular tone will follow. In addition, IDyOM estimates information content (*IC*) as the negative logarithm of the probability of a given tone:

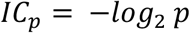

Thus, the more unexpected or surprising a tone is, the lower its probability and the larger its corresponding IC value. Note that mean IC can be used as an indirect measure of uncertainty, since less predictable contexts tend to have higher IC levels in the long run.

Both entropy and IC were used to estimate the predictability of the stimuli. For this, we first transformed the melodies to pitch-class representations in which the twelve tones of the musical scale system are coded without reference to the octave they belong to. This is consistent with the way Western tonal and atonal music are structured, and it allowed us to avoid pitch-alphabet discrepancies between the stimuli and the training corpora. Thus, pitch-class representations were used to train the model and predict pitch-class continuations in the melodies.

We estimated mean entropy and IC for each melody based on two training sets: a Western tonal corpus consisting of hymns and folk songs and an atonal corpus consisting of pieces by Hindemith and Varese as well as a set of atonal melodies created for a previous study (Dean and Pearce, 2016). These corpora constituted the long-term training of the model and simulated the knowledge of musical styles that a listener acquires during their lifetime. Here, we estimated separate values for each of the two training sets in order to cover the possibility that listeners’ expectations involved knowledge of tonal and atonal music.

The main IDyOM configuration we used is known as “*both*+”. This involves training the model with both a long-term corpus and the short-term regularities of the current stimuli, while at the same time incorporating the latter into the long-term component (as indicated by the “+” symbol). In addition, we obtained estimates in which no long-term knowledge was used—i.e., only short-term regularities were used. In this case, where estimates were driven by the statistics of the stimuli alone, the configuration is known as “*stm*”.

Figure 8 shows model estimates for the different conditions using the different corpora. For all three IDyOM configurations, mean entropy and IC values are higher for atonal than for tonal melodies. The difference in entropy was most pronounced in the no-corpus configuration, whereas the difference in IC was most pronounced in the tonal-corpus configuration. Furthermore, entropy was lowest in both conditions when the training corpus was tonal, followed by when the training corpus was atonal and no-corpus. This indicates that long-term knowledge of the Western tonal scale generally aids predictability, but the statistical properties of the stimuli themselves are sufficient to distinguish between the two conditions. This can be also appreciated in Figure 8*C*, where the distribution of pitch classes in atonal melodies is much more uniform than in tonal melodies, thus indicating higher entropy levels. In conclusion, IDyOM estimates corroborate higher uncertainty in atonal as compared with tonal stimuli.

### 4.3 MEG Procedures and Recordings

In the MEG session, participants were informed about the procedures both orally and in writing and gave their consent. Afterwards, they filled out a questionnaire with health-related and demographic questions. Four electrodes were applied on skin locations for EOG, two for ground and reference (on the forehead) and two for ECG (one each above the left and right collarbones). Once participants sat down in the MEG seat, three fiducial coils and in-ear headphones for the sound transmission were attached and the screen was positioned 50 cm in front of them. The MEG session started with a 10-minute resting-state recording during which participants had to look at a fixation cross set on a green background. Before beginning the task relevant to this study, participants carried out another paradigm in which they had to listen freely to excerpts from real music pieces for approximately 50 minutes. This paradigm was added to investigate naturalistic listening of musical pieces. The corresponding analyses and results will be published elsewhere. After a break in which participants were allowed to move a little and close their eyes for resting, the MMN paradigm started. They were instructed to watch a silent nature documentary without subtitles and were told to move as little as possible and pay no attention to the melodies. The MMN paradigm lasted approximately 20 minutes and was interrupted by one break between the tonal and atonal blocks. The MEG session ended with a tone localizer (lasting 6 minutes), where two different tones were played with random inter-onset intervals. After the electrodes were detached from their skin, participants answered in writing a few questions as to whether they felt sleepy during the MEG session, had any headache or back pain, and generally felt comfortable in the scanner. In a second session that took place on another day participants first underwent a structural MRI scan used for the source localization of the MEG sensor data inside the brain. Subsequently, they were asked to fill in a questionnaire about their musical backgrounds (general factor of the Goldsmiths Musical Sophistication Index; Schaal et al., 2014), the “Need for Cognition Scale” (Bless et al., 1994; Cacioppo et al., 1996) and the “Tellegen Absorption Scale” (Ritz and Dahme, 1995; Tellegen and Atkinson, 1974). Participants were then financially compensated for their participation.

MEG recordings were performed on a 275-channel whole-head MEG system with axial gradiometers (Omega 2000, CTF Systems Inc., Port Coquitlam, Canada) in a magnetically shielded booth. Data were acquired at a sampling rate of 1200 Hz, and online denoising (higher-order gradiometer balancing) was applied. Participants’ head positions relative to the MEG sensors were tracked continuously so that any head displacements could be corrected in the break by giving them instructions as to how to move back to their original positions. This was done by using the FieldTrip toolbox (http://fieldtrip.fcdonders.nl) (Stolk et al., 2013). T1-weighted MRIs were recorded with a 3 Tesla scanner (Siemens Magnetom Trio, Siemens, Erlangen, Germany).

### 4.4 MEG Analysis

#### 4.4.1 Preprocessing of MEG Data

To preprocess the data, the raw signal was first visually inspected with MNE-python (Gramfort et al., 2014) in order to check for deficient channels. None of the datasets showed irregular signals in any channel. The subsequent analyses were conducted with the FieldTrip toolbox (Oostenveld et al., 2011) in MATLAB (R2019b, The MathWorks, Natick, USA). Eye-blink movements and heartbeat artifacts were detected by means of a semi-automatic independent component analysis (ICA) routine that involved visual inspection for deciding which components to reject. In the next step, epochs between –100 ms and 400 ms around sound onset were extracted and baseline-corrected with a pre-stimulus baseline of 100 ms. Jump artifacts were automatically rejected if a trial exceeded a predefined z-value of 30. Both a low-pass filter (cut-off frequency of 40 Hz) and a high-pass filter (cut-off frequency of 0.6 Hz) were applied. The data were then downsampled to 250 Hz. Since 14 of the datasets were recorded with 271 and the remaining six datasets with only 270 sensors (one channel broke during the period of the data collection), corresponding missing channels were interpolated followed by the rejection of ICA components. Epochs corresponding to standard sounds— excluding those preceded by a deviant—and epochs corresponding to deviant sounds for each feature were averaged for each participant and condition. Finally, a new baseline correction was applied to the average evoked responses. MMNm difference waves were computed by subtracting the ERFs of standards from the ERFs of deviants.

#### 4.4.2 Statistical Analyses

For the first part of the statistical analyses, we used a mass-univariate approach in which two-sided dependent samples t-tests were performed. Cluster-based permutations (Maris and Oostenveld, 2007) were applied in order to correct for multiple comparisons. The cluster-forming significance threshold was .05, and the chosen statistic was the maximal sum of clustered T-values (maxsum). The number of iterations was 10,000. The tests were restricted to a window between 75 and 300 ms after sound onset (Näätänen et al., 2007).

To assess the presence of the MMNm, standards and deviants for each feature were contrasted for each condition and deviant feature separately. To assess the effect of tonal uncertainty, the MMNm difference waves were then compared between conditions. Since we performed a separate test for each deviant feature, a Bonferroni correction for multiple comparisons was applied by multiplying p-values by 4.

Further analyses were carried out to explore whether mean amplitudes and peak latencies of the MMNm were affected by tonal uncertainty. These analyses allowed us to test for feature-specific effects and more carefully assess main effects and interactions. For each participant, mean amplitudes and peak latencies were computed per feature, condition, and hemisphere. This was done by averaging the response ±25 ms around the peak, defined as the highest local maximum in the mean of a selection of channels in each hemisphere. These were the channels with the largest P50 response from a grand average of standard sounds across conditions, thus ensuring that the selected channels adequately captured auditory evoked activity while avoiding bias. Note that MMN auditory evoked fields exhibited a classical bipolar configuration (see Figure 2) with negative gradients in left anterior and right posterior channels and positive gradients in left posterior and right anterior channels (for a similar pattern, see magnetometer data in Quiroga-Martinez et al., 2020b). For this reason, we flipped the sign of activity in negative-gradient channels during peak finding. In the output, however, MMNm gradient amplitudes were given a negative sign for consistency with the negative polarity of the component in EEG.

Using R (RStudio, Version 1.3.959) and the lme4 library (Bates et al., 2015), several linear mixed effect models were calculated for each deviant feature with either mean amplitude or peak latency as the dependent variable. They were compared by means of likelihood ratio tests and built incrementally from intercept-only models by adding one factor at a time until a full model was reached. This included the relevant main factors and their two- and three-way interactions (see Table 1). Subject-wise random intercepts were included in all models. Since there were too few data points per condition, hemisphere, and feature, random slopes were not included in order to avoid overfitting. Models were also assessed with Akaike Information Criteria (AIC). Post-hoc, pairwise contrasts were also performed with the *emmeans* library (Lenth et al., 2019).

#### 4.4.3 Source Localization

For source localization, we used the Linearly Constrained Minimum Variance (LCMV) beamformer technique (Veen et al., 1997), as implemented in FieldTrip. Head models were obtained from individual MRI scans, which were manually realigned to match the head coordinate system defined by the MEG fiducial coils and were subsequently projected to a 7mm-resolution grid template in MNI coordinates provided by FieldTrip. Source localization was performed on the grid back-transformed into head coordinates, while group-level analyses were performed on source estimates projected onto the normalized MNI space. After tissue segmentation, leadfields were computed using a single shell volume conduction model (Nolte, 2003).

To compute the inverse solution with the LCMV beamformer technique, the covariance matrix for a particular deviant feature and condition was obtained from all trials representing standard and deviant sounds. The regularization parameter λ, which makes the solution more or less focal, was set to 10%. The orientation of each source was selected to be the one that maximizes the total power of source-level signal. The inverse solution was then used to estimate the standard, deviant, and difference (deviant-standard) source-level average power time-series for each voxel of the grid inside the brain.

Standard and deviant power time-series from all subjects were submitted to a two-sided paired-samples t-test to statistically test the presence of the MMNm in a cluster-based permutation approach, with 1000 iterations and p ≤ 0.05 as cluster-forming threshold. A significant positive cluster represents the case where the deviant has higher power than the standard, and a negative cluster represents the opposite. The interval of interest in which this statistical analysis was performed was selected to be –25 to +25 milliseconds around individual MMNm peaks. The grand average of the source power obtained from localizing the sensor-level difference between standard and deviants is displayed in Figure 5.

### 4.5 Behavioral Experiment

We conducted a separate online behavioral experiment on the platform www.pavlovia.org, using Psychopy (v2020.2.2). The sample comprised thirty-nine non-musicians (17 female, mean age = 36.77 ± 12.06, mean number of years of musical training = 0.48 ± 0.83) of whom none took part in the MEG experiment. They were selected from a larger pool of 59 non-musicians based on the inclusion criteria defined for the MEG experiment (no instrumental or singing training before the age of 10 and no such training for longer than three years). The remaining participants, albeit being non-musicians, did not fulfill the criteria. For transparency, in supplementary material 5, we report the analysis of the full sample. We used exactly the same melodies employed in the MEG experiment. The task required participants to answer, after each melody, whether there was a deviant note (yes=1/no=2) and how certain they were about their response (from 1 = “not certain at all” to 7 = “completely certain”). The target sounds were the same out-of-tune pitch deviants used in the MEG protocol. We decided to use only the pitch deviants since previous research showed that they were the most affected by contextual uncertainty. Furthermore, having included the other deviants would have significantly prolonged the online experiment which may have been detrimental for participants’ engagement and performance. The deviants were introduced in the second half of the melodies. There were 12 regular and 12 deviant trials per condition (tonal, atonal). This meant that each melody had two regular and two deviant versions, each one transposed to a different key or starting tone. Trial order was randomized for each participant. Participants were asked to use headphones to complete the task. The experiment lasted approximately 10 minutes and was approved by the internal review board (IRB) of the Danish Neuroscience Center at Aarhus University.

To analyze the data, we employed signal detection theory by calculating sensitivity (*d*’) and bias scores (criterion, or *c*) (Stanislaw and Todorov, 1999). We compared these measures between conditions with two-sided paired-samples *t*-tests. For confidence ratings, we employed ordinal logistic regression in the form of a mixed-effects cumulative-link model with subject-wise random intercepts, as implemented in the “ordinal” library in R (Christensen, 2019). The factors condition (tonal/atonal), deviance (regular melody/deviant melody), and their interaction were included as predictors. *p*-values were estimated for each parameter in the model through Wald tests. We hypothesized reduced sensitivity scores and lower confidence ratings in atonal as compared to tonal melodies.

## Supporting information

Supplementary material 2: Pitch content of the tonal and atonal condition.

Supplementary material 3: Peak coordinates in MNI space.

Supplementary material 4: Behavioral analysis.

Supplementary material 5: Behavioral analysis with full participant sample.

Supplementary material 1: Melodies of the atonal condition.

## CRediT authorship contribution statement

**Iris Mencke**: Conceptualization, Methodology, Formal analysis, Investigation, Writing - original draft, Visualization, Project administration, Funding acquisition. **David Quiroga-Martinez**: Conceptualization, Methodology, Formal analysis, Writing - original draft, Visualization. **Diana Omigie**: Conceptualization, Methodology, Writing – review & editing, Supervision. **Niels Trusbak Haumann**: Software, Writing – review & editing. **Georgios Michalareas**: Formal analysis, Writing – review & editing. **Franz Schwarzacher**: Formal analysis, Writing – review & editing. **Peter Vuust**: Conceptualization, Methodology, Writing – review & editing, Funding acquisition. **Elvira Brattico**: Conceptualization, Methodology, Writing – review & editing, Supervision, Funding acquistion.

## Acknowledgments

We thank William Martin for the profound english editing. Many thanks to Roger Dean who generously provided the atonal corpus for the computational measurements of our stimuli. We thank Simone Franz, Valerie Zipper, Franziska Debus, Daniela van Hinsberg and Jan Eggert for helping with the data collection. Thanks to Johanna Rimmele and Marina Kliucko for their constant support during the entire process. The Center for Music in the Brain is funded by the Danish National Research Foundation (DNRF 117). IM received funding from the Graduate School of Health (Aarhus University).

## Supplementary Material

Supplementary material 1: Melodies of the atonal condition.

Supplementary material 2: Pitch content of the tonal and atonal condition.

Supplementary material 3: Peak coordinates in MNI space.

Supplementary material 4: Behavioral analysis.

Supplementary material 5: Behavioral analysis with full participant sample.

