## Supplementary figures and images for "Prediction Under Uncertainty: Dissociating Sensory from Cognitive Expectations in Highly Uncertain Musical Contexts"

### Supplementary material 2: Pitch content of the tonal and atonal condition.

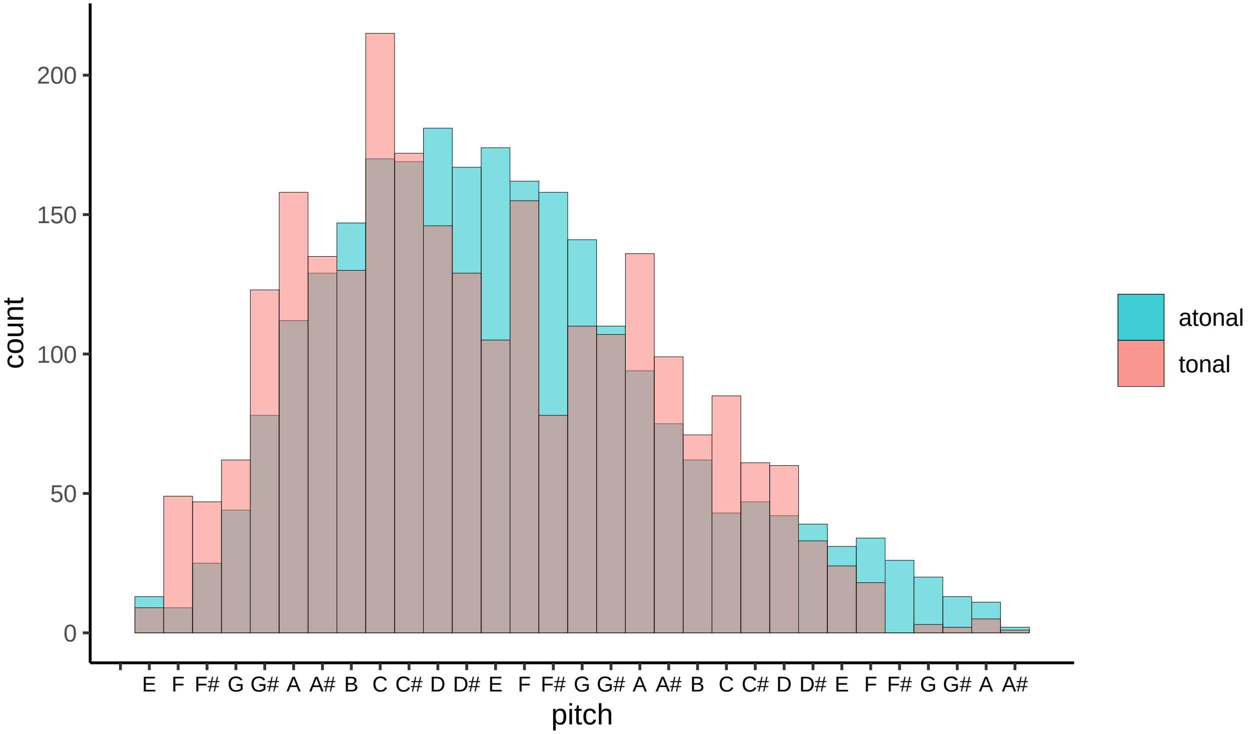
