## Supplementary material 3: Peak coordinates in MNI space. for "Prediction Under Uncertainty: Dissociating Sensory from Cognitive Expectations in Highly Uncertain Musical Contexts"

| condition | feature | hemisphere | x | y | z |
|---|---|---|---|---|---|
| tonal | pitch | right | 56 | -20 | 4 |
| tonal | pitch | left | -70 | -36 | 4 |
| tonal | intensity | right | 56 | -28 | 4 |
| tonal | intensity | left | -64 | -36 | 4 |
| tonal | timbre | right | 56 | -24 | -4 |
| tonal | timbre | left | -64 | -44 | 12 |
| tonal | location | right | 56 | -32 | 4 |
| tonal | location | left | -64 | -36 | 4 |
| atonal | pitch | right | 48 | -68 | -12 |
| atonal | pitch | left | -58 | -66 | 12 |
| atonal | intensity | right | 56 | -24 | -4 |
| atonal | intensity | left | -64 | -28 | 12 |
| atonal | timbre | right | 56 | -28 | 4 |
| atonal | timbre | left | -66 | -44 | 4 |
| atonal | location | right | 56 | -32 | 4 |
| atonal | location | left | -70 | -36 | 4 |
