## Supplementary material 4: Behavioral analysis. for "Prediction Under Uncertainty: Dissociating Sensory from Cognitive Expectations in Highly Uncertain Musical Contexts"

*David R. Quiroga-Martinez*

*5 feb 2021*

In this document, we show the results of an analysis including the 39 participants of the behavioral study who fulfilled the strict inclusion criteria.

Let's load the required libraries:

```
library(reshape2)
library(MASS)
library(lme4)
library(effects)
library(ggplot2)
library(viridis)
library(RColorBrewer)
library(ordinal)
library(cowplot)
library(kableExtra)
library(knitr)
library(broom)
library(broom.mixed)
library(dplyr)
```

Now, load the data:

```
d <- read.table('clean_data/dataset.csv', header = T, sep = ',') # load data

## To reproduce the original analyses with the stricter inclusion criteria
## (39 participants) uncomment these lines to select the subjects:

d <- d[d$prof < 2,] # self-declared as non-musicians
d <- d[d$yomt < 4,] # less than 4 years of musical training
d <- d[d$somt > 9|d$somt ==0,] # started musical traingin at 10 or older
```

After calculating d-prime scores (see Rmd script for the code), we run a t-test on them:

```
t.d <- t.test(d~cond, data= dprime, paired = T)
kable(tidy(t.d)%>%mutate_if(is.numeric,round,3))
```

| estimate | statistic | p.value | parameter | conf.low | conf.high | method | alternative |
|---|---|---|---|---|---|---|---|
| 0.646 | 5.15 | 0 | 38 | 0.392 | 0.9 | Paired t-test | two.sided |

The results show a clear effect of condition on d-prime scores. Now we run another t-test for criterion scores:

```
t.cr <- t.test(c~cond, data= dprime, paired = T)
kable(tidy(t.cr)%>%mutate_if(is.numeric,round,3))
```

| estimate | statistic | p.value | parameter | conf.low | conf.high | method | alternative |
|---|---|---|---|---|---|---|---|
| 0.067 | 0.692 | 0.493 | 38 | -0.129 | 0.262 | Paired t-test | two.sided |

The results show no significant effect of condition on criterion scores. Now we fit a (full) cummulative link mixed model of confidence ratings with the two factors and their interaction:

```r
conf <- clmm(as.factor(conf)~cond*deviance + (1|sub), data = d, link = "logit")
```

We inspect the parameters (odds ratios) of the model and the p-values:

```r
kable(tidy(conf,conf.int=T, conf.level = 0.95,exponentiate = T) %>%
        select(-c('std.error','coefficient_type')) %>%
        mutate_if(is.numeric,round,3))
```

| term | estimate | statistic | p.value | conf.low | conf.high |
|---|---|---|---|---|---|
| 1\|2 | 0.010 | -13.399 | 0.000 | 0.005 | 0.019 |
| 2\|3 | 0.044 | -9.379 | 0.000 | 0.023 | 0.085 |
| 3\|4 | 0.133 | -6.176 | 0.000 | 0.070 | 0.252 |
| 4\|5 | 0.281 | -3.906 | 0.000 | 0.149 | 0.531 |
| 5\|6 | 1.410 | 1.063 | 0.288 | 0.748 | 2.658 |
| 6\|7 | 6.775 | 5.867 | 0.000 | 3.576 | 12.839 |
| condatonal | 0.541 | -5.007 | 0.000 | 0.425 | 0.688 |
| condatonal:deviancestd | 0.776 | -1.493 | 0.136 | 0.556 | 1.083 |
| deviancestd | 0.874 | -1.112 | 0.266 | 0.689 | 1.108 |

The results show that participants were less confident in the atonal condition. There was not a significant condition*deviance interaction, but participants still tended to give particularly lower ratings to the standard melodies in the atonal condition.

Given this result, we explore whether a similar pattern can be seen for deviance detection. We therefore run a logistic regression on accuracy, with condition (tonal/atonal), deviance (standard/deviant) and their interaction as predictors.

```r
lr <- glmer(acc~cond*deviance+(1|sub),data=d,
            family=binomial(link = "logit"))

kable(tidy(lr,conf.int=T, conf.level = 0.95,exponentiate = T) %>%
        select(-c('std.error','effect','group')) %>%
        mutate_if(is.numeric,round,3))
```

| term | estimate | statistic | p.value | conf.low | conf.high |
|---|---|---|---|---|---|
| (Intercept) | 1.849 | 4.578 | 0.000 | 1.421 | 2.406 |
| condatonal | 0.729 | -2.282 | 0.023 | 0.556 | 0.956 |
| deviancestd | 2.723 | 6.336 | 0.000 | 1.998 | 3.713 |
| condatonal:deviancestd | 0.694 | -1.720 | 0.085 | 0.458 | 1.052 |
| sd___(Intercept) | 0.561 | NA | NA | NA | NA |

Notably, the same (non-significant) trend can be seen here. Participants tended to be less accurate in atonal melodies, and more so when the melody was a standard. Nevertheless, note that accuracy was already lower for deviant melodies.

Finally, we make the corresponding plots (se Rmd script for the full code)

```r
plots <- align_plots(d.plot,c.plot,conf.plot2,conf.plot1,align = 'v',axis ='1')
joint_plots <- plot_grid(plots[[1]],plots[[2]],plots[[3]],plots[[4]],
                         labels = c('a','b','c','d'), ncol =2,nrow =2)
ggsave("joint_plots.png", plot=joint_plots,width = 180, height = 180,
       units = 'mm', dpi = 600)
joint_plots
```

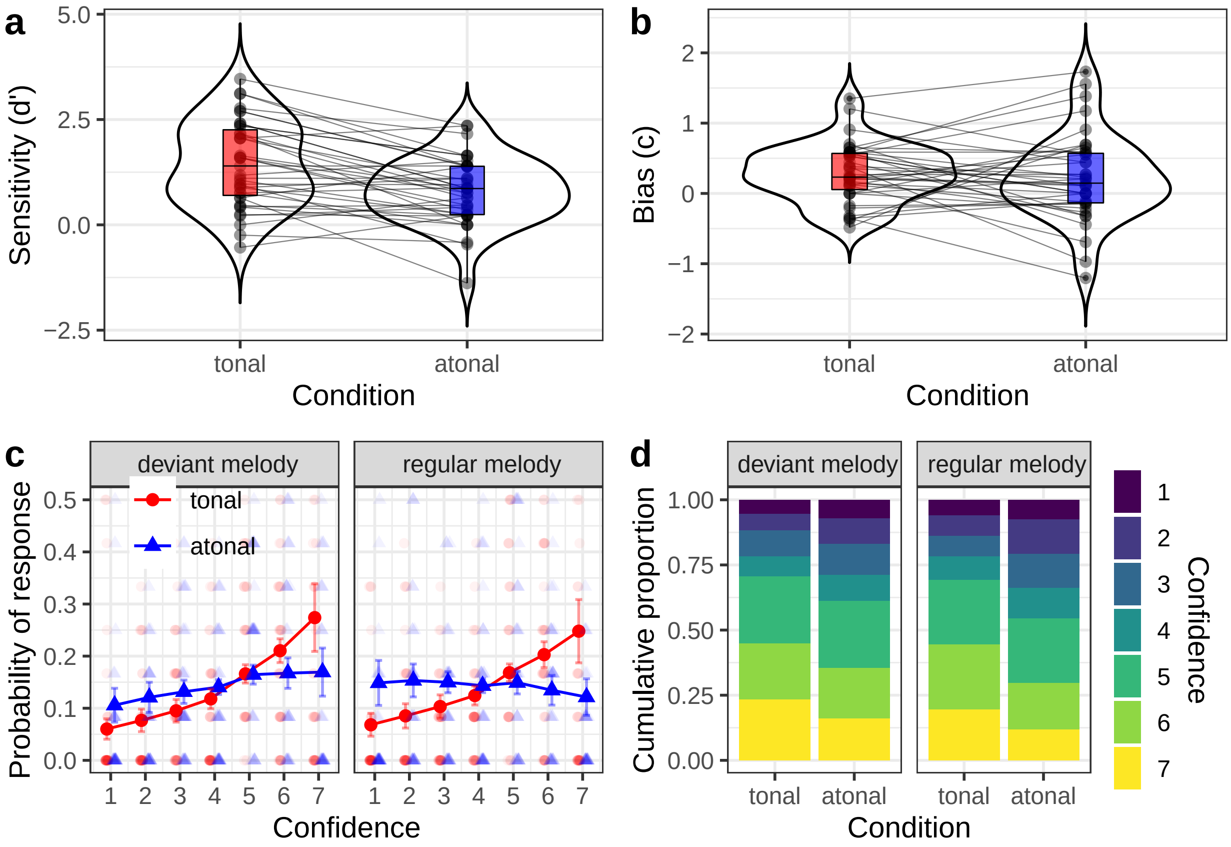
a
5.0
2.5
0.0
−2.5
Sensitivity (d')
tonal
atonal
Condition
b
2
1
0
−1
−2
Bias (c)
tonal
atonal
Condition
c
deviant melody
regular melody
tonal
atonal
0.5
0.4
0.3
0.2
0.1
0.0
Probability of response
1 2 3 4 5 6 7
1 2 3 4 5 6 7
Confidence
d
deviant melody
regular melody
1.00
0.75
0.50
0.25
0.00
Cumulative proportion
tonal
atonal
tonal
atonal
Condition
Confidence
1
2
3
4
5
6
7
