## Supplementary material 5: Behavioral analysis with full participant sample. for "Prediction Under Uncertainty: Dissociating Sensory from Cognitive Expectations in Highly Uncertain Musical Contexts"

### extended behavioral analysis

*David R. Quiroga-Martinez*

*5 feb 2021*

In this document, we show the results of a full analysis including all participants of the behavioral study.

Let's load the required libraries:

```
library(reshape2)
library(MASS)
library(lme4)
library(effects)
library(ggplot2)
library(viridis)
library(RColorBrewer)
library(ordinal)
library(cowplot)
library(kableExtra)
library(knitr)
library(broom)
library(broom.mixed)
library(dplyr)
```

Now, load the data:

```
d <- read.table('clean_data/dataset.csv', header = T, sep = ',') # load data

## To reproduce the original analyses with the stricter inclusion criteria
## (39 participants) uncomment these lines to select the subjects:

# d <- d[d$prof < 2,] # self-declared as non-musicians
# d <- d[d$yomt < 4,] # less than 4 years of musical training
# d <- d[d$somt > 9|d$somt ==0,] # started musical traingin at 10 or older
```

After calculating d-prime scores (see Rmd script for the code), we run a t-test on them:

```
t.d <- t.test(d~cond, data= dprime, paired = T)
kable(tidy(t.d)%>%mutate_if(is.numeric,round,3))
```

| estimate | statistic | p.value | parameter | conf.low | conf.high | method | alternative |
|---|---|---|---|---|---|---|---|
| 0.563 | 5.163 | 0 | 58 | 0.345 | 0.781 | Paired t-test | two.sided |

The results show a clear effect of condition on d-prime scores. Now we run another t-test for criterion scores:

```
t.cr <- t.test(c~cond, data= dprime, paired = T)
kable(tidy(t.cr)%>%mutate_if(is.numeric,round,3))
```

| estimate | statistic | p.value | parameter | conf.low | conf.high | method | alternative |
|---|---|---|---|---|---|---|---|
| 0.06 | 0.785 | 0.436 | 58 | -0.093 | 0.213 | Paired t-test | two.sided |

The results show no significant effect of condition on criterion scores. Now we fit a (full) cummulative link mixed model of confidence ratings with the two factors and their interaction:

```
conf <- clmm(as.factor(conf)~cond*deviance + (1|sub), data = d, link = "logit")
```

We inspect the parameters (odds ratios) of the model and the p-values:

```
kable(tidy(conf,conf.int=T, conf.level = 0.95,exponentiate = T) %>%
        select(-c('std.error','coefficient_type')) %>%
        mutate_if(is.numeric,round,3))
```

| term | estimate | statistic | p.value | conf.low | conf.high |
|---|---|---|---|---|---|
| 1\|2 | 0.008 | -16.064 | 0.000 | 0.005 | 0.015 |
| 2\|3 | 0.036 | -11.624 | 0.000 | 0.020 | 0.062 |
| 3\|4 | 0.110 | -7.826 | 0.000 | 0.063 | 0.191 |
| 4\|5 | 0.273 | -4.634 | 0.000 | 0.157 | 0.472 |
| 5\|6 | 1.296 | 0.930 | 0.353 | 0.750 | 2.241 |
| 6\|7 | 6.050 | 6.402 | 0.000 | 3.487 | 10.497 |
| condatonal | 0.511 | -6.706 | 0.000 | 0.420 | 0.622 |
| condatonal:deviancestd | 0.759 | -1.981 | 0.048 | 0.578 | 0.997 |
| deviancestd | 0.909 | -0.961 | 0.337 | 0.748 | 1.104 |

The results show that participants were less confident in the atonal condition. There was a condition*deviance interaction meaning that participants gave particularly lower ratings to the standard melodies in the atonal condition.

Given this result, now we explore whether the same interaction can be seen for deviance detection. We therefore run a logistic regression on accuracy, with condition (tonal/atonal), deviance (standard/deviant) and their interaction as predictors.

```
lr <- glmer(acc~cond*deviance+(1|sub),data=d,
            family=binomial(link = "logit"))

kable(tidy(lr,conf.int=T, conf.level = 0.95,exponentiate = T) %>%
        select(-c('std.error','effect','group')) %>%
        mutate_if(is.numeric,round,3))
```

| term | estimate | statistic | p.value | conf.low | conf.high |
|---|---|---|---|---|---|
| (Intercept) | 1.718 | 4.795 | 0.000 | 1.377 | 2.144 |
| condatonal | 0.757 | -2.474 | 0.013 | 0.607 | 0.944 |
| deviancestd | 2.833 | 8.163 | 0.000 | 2.207 | 3.638 |
| condatonal:deviancestd | 0.713 | -1.968 | 0.049 | 0.508 | 0.999 |
| sd___(Intercept) | 0.606 | NA | NA | NA | NA |

Notably, the same interaction can be seen here. Participants were less accurate in atonal melodies, and more so when the melody was a standard. Nevertheless, note that accuracy was already lower for deviant melodies.

Finally, we make the corresponding plots (se Rmd script for the full code)

```
plots <- align_plots(d.plot,c.plot,conf.plot2,conf.plot1,align = 'v',axis ='1')
joint_plots <- plot_grid(plots[[1]],plots[[2]],plots[[3]],plots[[4]],
                         labels = c('a','b','c','d'), ncol =2,nrow =2)
ggsave("joint_plots_extended.png", plot=joint_plots,width = 180, height = 180,
       units = 'mm', dpi = 600)
joint_plots
```

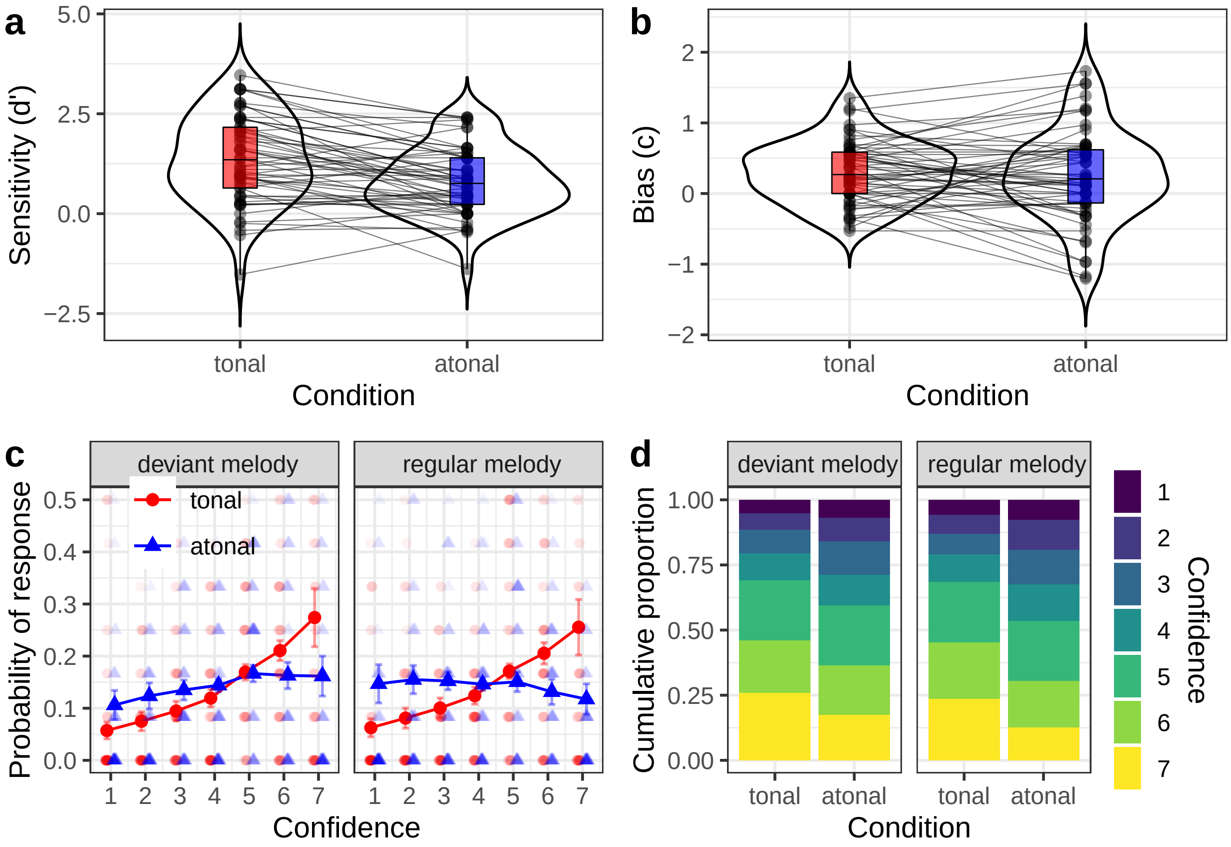

a
Sensitivity (d')
5.0
2.5
0.0
−2.5
tonal
atonal
Condition
b
Bias (c)
2
1
0
−1
−2
tonal
atonal
Condition
c
deviant melody
regular melody
tonal
atonal
Probability of response
0.5
0.4
0.3
0.2
0.1
0.0
1 2 3 4 5 6 7
Confidence
d
deviant melody
regular melody
Cumulative proportion
1.00
0.75
0.50
0.25
0.00
tonal atonal
tonal atonal
Condition
Confidence
1
2
3
4
5
6
7
