## Supplementary material 1: Melodies of the atonal condition. for "Prediction Under Uncertainty: Dissociating Sensory from Cognitive Expectations in Highly Uncertain Musical Contexts"

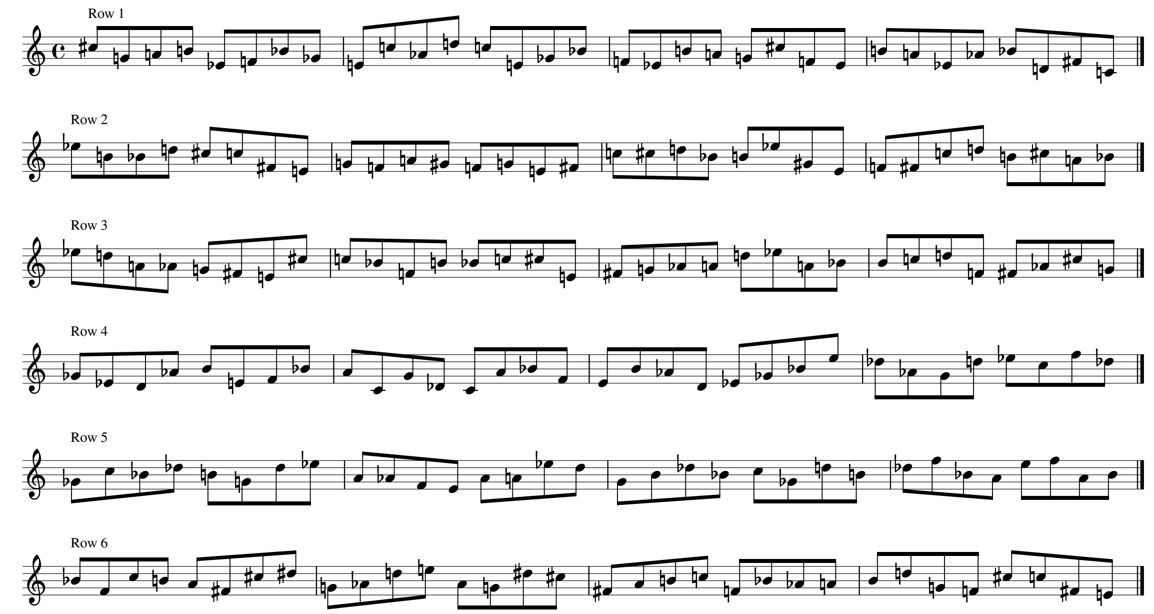


Six resulting melodies built on the basis of twelve-tone rows. They were used to generate the melodic material for the atonal condition.

References:

| | |
|---|---|
| Row 1 | Arnold Schönberg: Three Songs for Voice and Piano, Op. 48, „Mädchenlied“ |
| Row 2 | Anton Webern: Variationen for Piano, Op.27, Variation IV |
| Row 3 | Pierre Boulez: Structure 1a (row from Olivier Messiaens *Mode de Valeurs et de intensites*) |
| Row 4 | Arnold Schönberg, String Quartet No.3 , Op. 30 |
| Row 5 | Arnold Schönberg, Variations for Orchestra, Op. 31, Reihe 1b |
| Row 6 | Arnold Schönberg, Piano Piece, Op. 33a |
